# Cross-linked natural IntegroPectin films from Citrus biowaste with intrinsic antimicrobial activity

**DOI:** 10.1101/2022.01.28.478153

**Authors:** Elena Piacenza, Alessandro Presentato, Rosa Alduina, Antonino Scurria, Mario Pagliaro, Lorenzo Albanese, Francesco Meneguzzo, Rosaria Ciriminna, Delia F. Chillura Martino

## Abstract

Pectin recovered via hydrodynamic cavitation (IntegroPectin) from lemon and grapefruit agri-food waste intrinsically containing antimicrobial bioactive substances (flavonoids, phenolic acids, terpenes, and terpenoids) was used to generate innovative and eco-compatible films that efficiently inhibit the growth of Gram-negative pathogens. Extensive characterization of films confirmed the presence of these substances, which differently interact with the polysaccharide polymer (pectin), plasticizer (glycerol), surfactant (Tween 60), and cross-linker (Ca^2+^), conferring to these films a unique structure. Besides, IntegroPectin-based films constitute sustained, controlled, and slow-release (up to 72h) systems for bioactive substances in an aqueous environment. This feature is crucial for the good *in vitro* antimicrobial activity exerted by IntegroPectin films against three Gram-negative bacteria (two indicator pathogen strains *Pseudomonas aeruginosa* ATCC 10145, *P. aeruginosa* PAO1, and the clinical isolate *Klebsiella pneumoniae*) that are involved in the global emergence of the antimicrobial resistance.

## 1. Introduction

In the last ten years, the worldwide production of plastic and plastic-based materials increased up to 20-fold, generating large amounts of waste that excessively accumulate in the environment (Gouveia, Biernacki, Castro, Goncalves, & Souza, 2019; Shivangi, Dorairaj, Negi, Shetty, & 2021). This event poses crucial ecological and toxicological problems, as several synthetic polymers used for producing plastic-based materials can cause severe health issues and can be carrier of microorganisms (Shivangi et al., 2021; Gao, He, Sun, He, & Zeng 2019; Sucato et al., 2021). Thus, we are witnessing an increased demand for safer alternatives that can replace non-biodegradable plastic-based materials. Biopolymers can represent bio- and eco-friendly replacements for plastic ones, being biodegradable, abundant, renewable, and, in most cases, even edible (Gomaa, Fawxy, Hifney, & Abdel-Gawad, 2018). Among the plethora of plastic materials that biopolymers can substitute, films are one of the most studied and explored, as they can find applications in a broad spectrum of fields (Gouveia et al., 2019; Mishra, Banthia, & Majeed, 2012; Martau, Mihai, & Vodnar, 2019; Zhu, 2021). Some of the most used biopolymers for film development are polysaccharides (pectin, cellulose, starch, sodium alginate, chitosan, and gums) due to their easy processing and cost-effectiveness (Nogueira, de Oliveira, Velasco, & Fakhouri, 2020). The different polysaccharide features (*e.g.*, molecular weight, composition, conformation, glycosidic bonds, and functional groups) confer diverse physical-chemical properties to benefit for developing a wide array of bio- and eco-compatible films (Zhu, 2021). For instance, polysaccharide films can act as efficient barriers against oxygen, carbon dioxide, and lipids (Gouveia et al., 2019; Nisar et al., 2019), or anticancer and antidiabetic agents (Zhu, 2021).

Pectin is a branched and anionic polysaccharide widely used in food, medical, and pharmaceutical industries that can be recovered from biological resources or waste, such as citrus peel or apple pomace (Fidalgo et al., 2016; Yu, Shen, Song, & Xie, 2018; Meneguzzo et al., 2019). Pectin’s structure constitutes of 1,4-D-galacturonic acid units with partially or fully methyl esterified carboxylic groups (Presentato et al., 2020a; Shivangi et al., 2021). Key parameters influencing both rheological and biological (antimicrobial, antioxidant, anti-inflammatory, anticancer) properties of pectin are the degree of pectin esterification, its molecular weight, and the amount of homogalacturonan (HG) smooth and RG-I “hairy” regions (Yu et al., 2018), which make pectin a highly versatile choice over synthetic polymers. Furthermore, pectin is an excellent candidate for generating films due to its partially crystalline structure and polyelectrolyte nature (Nisar et al., 2019; Balik, Argin, Lagaron, & Torres-Giner, 2019). However, the brittleness, intrinsic hydrophilicity, and low mechanical strength of neat pectin films highlight disadvantages for their applications (Gouveia et al., 2019; Nisar et al., 2019; Balik et al., 2019). In this regard, plasticizers such as polyols (*e.g*., glycerol or sorbitol) are generally added to pectin formulations to enhance the motion of polysaccharide chains and reduce their intra- and inter-molecular forces, improving the mechanical integrity and flexibility of pectin-based films (Balik et al., 2019). Similarly, *in situ* cross-linking with metal cations (Ca^2+^, Zn^2+^ or Mg^2+^) can lower the pectin solubility in water, simultaneously creating a three-dimensional matrix where metal cations can localize within the twisted HG chains, improving physical-chemical properties of the films (Gao et al., 2019; Balik et al., 2019). Besides, pectin, being an anionic polymer, is suitable for integrating and carrying phytochemical and bioactive compounds (polyphenols, flavonoids, essential oils). These substances can enhance films’ properties and confer bio-applicative features, such as antioxidant, antimicrobial, and anticancer activities (Mishra et al., 2012; Nogueira et al., 2020; Nisar et al., 2019).

Pectins extracted by hydrodynamic cavitation (HC) from waste lemon and grapefruit peels (henceforth indicated as lemon -LIP- and grapefruit IntegroPectin -GIP-) revealed their uniqueness over the commercial citrus pectin (CP). This phenomenon relies on the presence, in the former, of polyphenols and terpenes (Presentato et al., 2020a; Scurria et al., 2021a; Scurria et al., 2021b) that provide a fundamental economic and practical value compared to films that need to be functionalized or loaded with these compounds. Indeed, bioactive substances within IPs convey to the latter biological and chemical properties that are highly relevant for their application, as they can act as antimicrobial, antioxidant, neuroprotective, and antiproliferative agents without exerting cytotoxic effects on human cell lines (Presentato et al., 2020a; Nuzzo et al., 2021). Specifically, LIP and GIP exhibited strong *in vitro* antimicrobial activity against Gram-negative and -positive opportunistic pathogen indicator strains (*i.e., Staphylococcus aureus* and *Pseudomonas aeruginosa*) that we find in almost every environmental niche (Presentato et al., 2020a; Presentato et al., 2020b). This evidence is of paramount interest and importance, as IPs can constitute innovative antimicrobials to counteract the health issue related to Antimicrobial Resistance (AMR) in microorganisms, which, according to current projections, will be responsible for more than ten million deaths and an economic toll comparable to the 2008 financial crisis by 2050 (Wellcome Trust, 2016). Thus, research on new, innovative, and biocompatible antimicrobials targeting microbial resistance mechanisms is nowadays flourishing. For instance, natural substances (plant-derived flavonoids, polyphenols, and phenolic acids) or metals, such as silver in colloidal form, have been widely investigated for their antimicrobial potential (Russo et al., 2016; Wright, 2017; Biharee, Sharma, Kumar, & Jaitak, 2020; Ciriminna, Albo, & Pagliaro, 2020). Yet, the extensive use and exposure of pathogenic microorganisms to silver causes the emergence of bacterial resistance mechanisms against this metal (Panacek et al., 2018; Hosny, Rasmy, Abdoul-Magd, Kashef, & El-Bazza, 2019), leading researchers to investigate efficient, eco-compatible, and innovative alternatives.

In the present study, we explored the possibility of using LIP and GIP formulations to generate bio-, eco-compatible and antimicrobial films. Films’ structure and nature were investigated through Fourier Transform Infrared spectroscopy in Attenuated Total Reflectance (ATR-FTIR) mode, solidstate ^13^C Cross Polarization Magic-Angle Spinning Nuclear Magnetic Resonance (CP MAS NMR), and X-ray Diffraction (XRD). Moreover, the potentiality of IP films (IPFs) in releasing bioactive substances with antimicrobial value in an aqueous environment was assessed for up to 72h. The antimicrobial potential of IPFs was evaluated against three Gram-negative strains (two pathogen indicator strains *P. aeruginosa* ATCC 10145 and *P. aeruginosa* PAO1 and a clinical isolate of *K. pneumoniae*), which, being more tolerant and resistant towards antibiotics than Gram-positive ones (Breijyeh, Jubeh, & Karaman, 2020), result more difficult to defeat.

## 2. Experimental Section

### 2.1. Materials

All the reagents were purchased from Sigma-Aldrich^®^ (Milan, Italy), except for calcium chloride (CaCl_2_) that was purchased from Carlo Erba Reagenti (Rodano, Italy). LIP and GIP were obtained through HC of waste citrus peels deriving from organically grown lemons and grapefruits by a citrus processing factory located in Sicily (Campisi Citrus, Siracusa, Italy).

### 2.2. Methods

#### 2.1.1. Preparation of pectin-based films

1 g of LIP, GIP, or CP (DE = 69%, galacturonic acid >74.0%, dried basis) was dissolved in 50 g distilled water obtained with a Thermo Scientific Barnstead Smart2Pure water purification system (Thermo Fisher Scientific, Waltham, Massachusetts, USA) at room temperature. IP samples readily dissolved, whereas the dissolution of CP required prolonged stirring and heating at 50-60°C. 800 mg of glycerol (99.5% pure) and 100 mg of Tween 60 were added to each pectin dispersion. The resulting mixtures were emulsified at 6,000 rpm for 3 min with an Ultra-Turrax T 25 (IKA-Werke, Staufen, Germany), followed by addition of a 5 ml aliquot of a 1% (w/v) solution of CaCl_2_ anhydrous under vigorous stirring. Finally, dispersions were casted onto Petri dishes (8 cm in diameter), and CP, LIP, and GIP films (henceforth named as CPF, LIPF, and GIPF, respectively) were obtained by solvent casting in a UF 30 oven (Memmert, Schwabach, Germany) at 40°C for 24 h.

#### 2.2.2. Physical-chemical characterization of pectin-based films

Pectin-based films were characterized through ATR-FTIR and ^13^C CPMAS NMR spectroscopies alongside XRD analysis.

ATR-FTIR spectra were collected, by using an FTIR Bruker Vertex Advanced Research Fourier Transform Infrared spectrometer (Bruker, Billerica, MA, USA) equipped with a Platinum ATR and a diamond crystal, in the 70-4000 cm^-1^ range (later resolution of 2 cm^-1^ and 200 scans). The Degree of Esterification (DE) of CPF, LIPF, and GIPF was determined following the Fidalgo-Ilharco equation (Fidalgo et al., 2016):

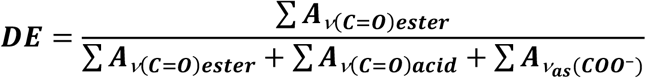

where A indicates the integrated area obtained after the spectra deconvolution by non-linear leastsquares fitting of the 1800-1400 cm^-1^ region for both the analyzed samples.

^13^C CPMAS NMR was performed by using a Bruker Advance II 400 (9.4T) spectrometer Bruker BioSpin GmbH, Rheinstetten, Germany) operating at 400.15 MHz and 100.61 MHz for ^1^H and ^13^C nuclides, respectively. Fifty milligrams of each film were packed in 4 mm diameter zirconia rotors sealed with KEL-F caps. The spectra were acquired at 300 K and a rotation frequency of the magic angle of 6 kHz by setting a 90° impulse on ^1^H of 4.5 ms, a delay time of 3 s, and 400 scans. An adamantane standard sample was exploited as standard for both the optimization of the Hartman-Hahn condition and the chemical shift reference (29.5 and 38.6 ppm).

XRD patterns were collected through a Brucker ecoD8 ADVANCE (Bruker, Billerica, MA, USA) spectrometer working in the θ-2θ geometry equipped with a Cobalt tube (Co; λ = 1.79 Å) and a LYNEXEYE (1D mode) detector operating at 40 kV and 25 mA. XRD patterns were recorded in the 5.00°-70.0° (2θ) range with a 1 mm window, an increment of 0.0198°, and time *per* step of 1.00 s.

FTIR and ^13^C CPMAS NMR spectra, and XRD patterns were analyzed by using OPUS(7.5), MestReNova(11.0.0), and Match!3 software, respectively. All the graphical representations and spectral deconvolutions were obtained through OriginPro^®^ software (OriginLab Corporation, Northampton, MA, USA).

#### 2.2.3 Release kinetics of active biomolecules by IPFs

IPFs’ capability of releasing bioactive substances was evaluated over time (2-72h) through UV-Visible spectroscopy. Sections (1 cm x 1 cm) of pectin-based films were cut and transferred into microtubes containing 2 ml of Tryptic Soy Broth (TSB) medium, which were incubated at room temperature. Aliquots (50 μl added to 1.5 ml of distilled water) of TSB medium were collected after 2, 4, 6, 8, 10, 24, 36, 48, and 72h of film incubation and their absorption spectra were recorded in the 200-700 nm range through a Beckman DU 800 spectrophotometer (Beckman Coulter Life Sciences, Milan, Italy). The experiments were performed in triplicate (*n* = 3) and absorption spectra were analyzed through OriginPro^®^ software.

Putative release kinetic profiles of bioactive substances from IPFs were obtained focusing on absorbance contributions that were consistent over time (LIPF: 270-275 nm, 306-315 nm, 340-350 nm; GIPF: 280-286 nm). Cumulative data were fitted to diverse kinetic models (*i.e.*, zero-order, first-order, second-order kinetics, Higuchi, Hixson-Crowell, and Ritger-Peppas models) (Bruschi, 2015) to study the mechanism(s) of bioactive substance release from IPFs over the timeframe considered.

#### 2.2.4 Antimicrobial activity of pectin-based films

The antimicrobial potential of LIPF, GIPF, and CPF was tested against indicator pathogen strains *P. aeruginosa* ATCC 10145, *P. aeruginosa* PAO1, alongiside the clinical isolate *K. pneumoniae*, as described elsewhere (Presentato et al., 2020a) with slight modifications. Briefly, a single colony of bacterial strains was pre-cultivated (16h) at 37 °C with shacking (160 rpm) in the TSB medium. The same medium was solidified by adding 15 g/L of bacteriological agar when needed. Bacterial cells were then inoculated [0.05% v/v corresponding to ca. 5×10^5^ colony forming units (CFU) ml^-1^] in the presence of 1 cm x 1 cm of LIPF, GIPF, or CPF and challenged for 24h according to the same pre-culturing conditions. The number of viable CFU ml^-1^ surviving the challenge exerted by pectin-based films was evaluated through the spot plate count method and compared to unchallenged cultures. The data are reported as average values (*n* = 3) of the CFU ml^-1^ in the logarithmic scale with standard deviation. Besides, LIPF and GIPF were tested for their ability to inhibit the growth of adherent bacterial cells on a solid medium. Specifically, ca. 1×10^7^ CFU ml^-1^ were plated onto TSB agar plates, allowing for a confluent growth, and IPFs were placed on top of the bacterial layer. Plates were then incubated at 37°C under static conditions for 24h. The subsequent day, inhibitions halos of the bacterial growth were qualitatively evaluated.

## 3. Results and Discussion

### 3.1. Physical-chemical characterization of commercial pectin- and IntegroPectin-based films

#### 3.1.1 ATR-FTIR spectroscopy

ATR-FTIR spectra of CPF, LIPF, and GIPF revealed vibrational modes attributable to the presence of pectin alongside bioactive substances, such as polyphenols, flavonoids, and terpenes (**Figure 1**; **Table S1**), being in line with previous observations reported for powder formulations (Presentato et al., 2020a; Scurria et al., 2021a; Scurria et al., 2021b).

**Figure 1.**
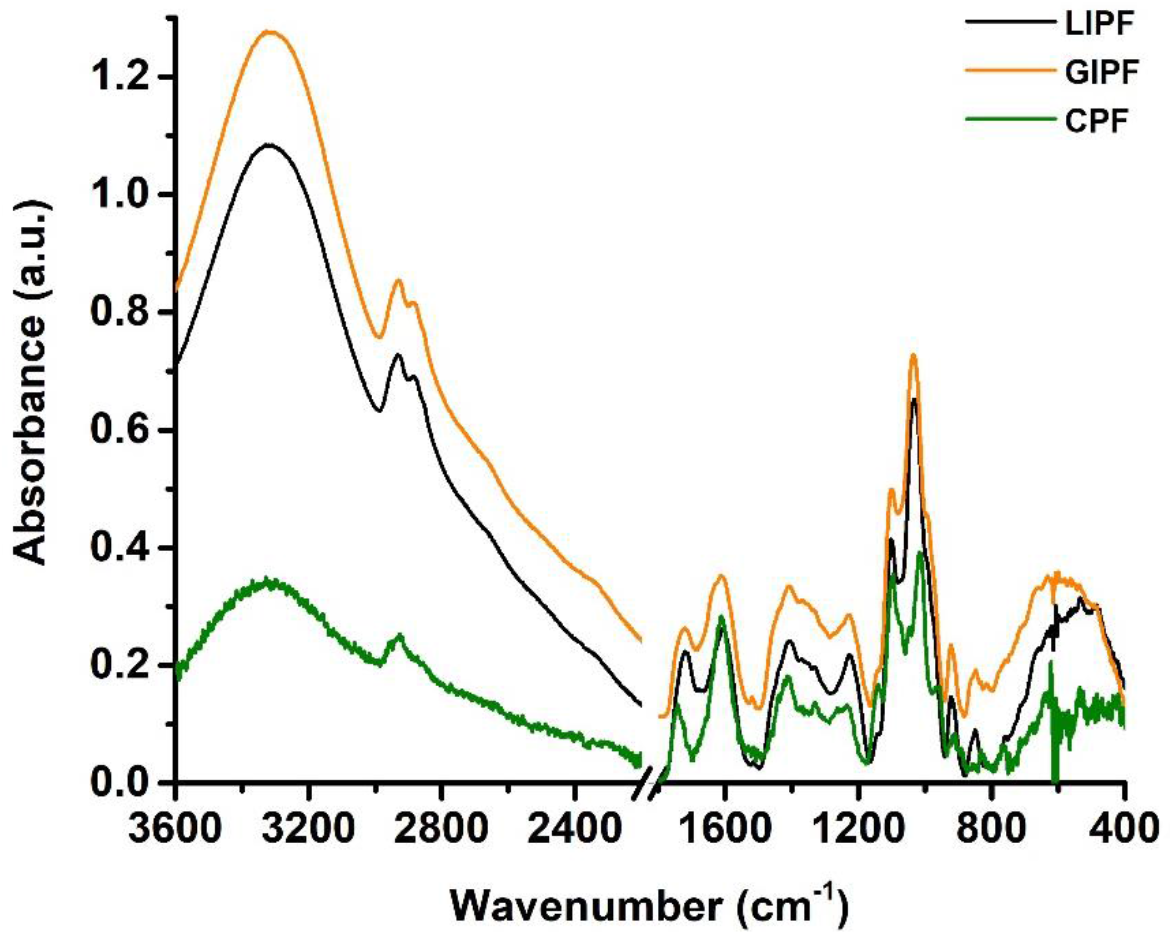
ATR-FTIR spectra collected for CPF, LIPF, and GIPF. For clarity, spectra were offset by 0.1 arbitrary units (a.u.).

Nevertheless, the filming process led to more intense IR absorption bands corresponding to -OH (ca. 3300 cm^-1^) and -CH_x_ (2930-2850 cm^-1^) stretching (Figure 1) in IPFs than powder forms (Presentato et al., 2020a), which partially derived from the addition of glycerol and Tween 60 as plasticizer and surfactant, respectively. The intense -OH stretching signal observed at ca. 3290 cm^-1^ for LIPF and GIPF also related to the presence of -OH rich biomolecules, such as polyphenols and flavonoids, that were instead absent in CPF, as indicated by the less pronounced and shifted (ca. 3320 cm^-1^) -OH stretching vibration (Figure 1; Table S1). The difference in intensity and shape of - OH vibration observed for IPFs over CPF can be caused by the development of intermolecular hydrogen bonds between either glycerol or Tween 60 with bioactive substances, which generally occur between carboxylic acid (for phenolic acids) or carbonyl (for flavonoids) moieties and -OH groups of glycerol or Tween 60 (Huaman-Castilla, Mariotti-Celis, Martinez-Ficuentes, & Perez-Correa, 2020).

The 1800-1300 cm^-1^ region of ATR-FTIR spectra featured IR absorption mostly attributable to - C=O (1750-1600 cm^-1^) and pectin ring (1400-1300 cm^-1^) vibration and deformation modes (Fidalgo et al., 2016; Presentato et al., 2020a), yet contributions deriving from phenolic acids, polyphenols, terpenes, and flavonoids were detected in LIPF and GIPF (Figure 1; Table S1). The -C=O stretching vibration of esterified carboxylic groups (ca. 1740 cm^-1^) was distinguishable only for CPF (Table S1), while the same contribution was absent, as maximum, for IPFs, likely due to its partial overlapping with the symmetric -C=O stretching vibration of carboxylate and nonconjugated keto groups (ca. 1715 cm^-1^) of bioactive substances (Lóránd, Deli, Molnár, & Toth, 2002; Heredia-Guerrero et al., 2014; Ricci, Olejar, Parpinello, Kilmartin, & Versari, 2015). More insights on IPFs’ structure and composition were obtained through a deconvolution of ATR-FTIR spectra in the 1800-1480 cm^-1^ region (**Figure S1**; **Table 1**).

**Table 1.** Deconvolution of LIPF and GIPF ATR-FTIR spectra by non-linear least squares fitting of the 1800-1480 cm^-1^ region.

| LIPF |  |  | GIPF |  |  | Vibrational mode<sup>a</sup> |
| --- | --- | --- | --- | --- | --- | --- |
| $\tilde{\nu}$ [cm<sup>-1</sup>]<sup>b</sup> | w<sup>c</sup> | A<sup>d</sup> | $\tilde{\nu}$ [cm<sup>-1</sup>] | w | A | |
| 1750 | 21.8 | 1.47 | 1750 | 21.0 | 1.28 | $\nu^e$ (C=O)<sub>ester</sub> |
| 1723 | 37.2 | 6.62 | 1724 | 38.9 | 5.74 | $\nu_s^f$ (C=O)<sub>carboxylate and nonconj. keto groups</sub> |
| 1676 | 85.6 | 14.40 | 1672 | 69.3 | 8.54 | $\nu$ (C=O)<sub>acid</sub> |
| | | | 1640 | 17.8 | 0.51 | $\delta^g$ (H<sub>2</sub>O); $\nu$ (C=C)<sub>uracyl</sub>; $\nu$ (C-C)<sub>phenyl</sub> |
| 1607 | 59.3 | 16.27 | 1604 | 64.8 | 18.23 | $\nu_{as}^h$ (COO<sup>-</sup>)<sub>polygalacturonic acid</sub>; $\nu$ (C-C) |
| 1513 | 22.9 | 0.423 | 1512 | 31.8 | 0.985 | $\delta$ (CH<sub>2</sub>) |

<sup>a)</sup> references are enlisted in the Supplementary Material; <sup>b)</sup> $\tilde{\nu}$ [ $\text{cm}^{-1}$ ] = wavenumber; <sup>c)</sup> $w$ = width; <sup>d)</sup> $A$ = integrated area; <sup>e)</sup> $\nu$ = stretching vibration; <sup>f)</sup> $s$ = symmetric vibration; <sup>g)</sup> $\delta$ = bending vibration; <sup>h)</sup> $as$ = asymmetric vibration.

In line with previous findings, IR bands of esterified carboxylic groups of pectin galacturonic acid (GalA) residues arose for IPFs at ca. 1750 cm^-1^ (Fidalgo et al., 2016; Wang et al., 2016; Bichara et al., 2016; La Cava, Gerbino, Sgroppo, & Gomez-Zavaglia, 2018) (Figure S1; Table 1). The width and area of these contributions were comparable between LIPF and GIPF (Table 1), alongside those observed for IP powders (Presentato et al., 2020a), suggesting a similar content of GalA for all formulations. IPFs featured also IR bands deriving from carboxylate and nonconjugated keto groups (ca. 1723 cm^-1^), nonesterified hydrogenated acidic carbonyl and conjugated keto groups, and carboxylic acid groups with strong H bonds (ca. 1676 cm^-1^) (Fidalgo et al., 2016) (Figure S1; Table 1). A higher contribution of -C=O vibration of acid groups was detected for LIPF than GIPF (Table 1), confirming the more acidic nature of the former previously suggested for IP powders (Presentato et al., 2020a). On the opposite, GIPF showed vibrational modes (1640 cm^-1^; Table 1) typical of uracyl and phenyl moieties of the most abundant flavonoids (*e.g*., naringenin and naringin) found in grapefruit (Scurria et al., 2021b), which were preserved by the HC extraction procedure (La Cava et al., 2018). Nevertheless, this contribution was noticeably lower, in terms of width and area (Table 1), than GIP powder (Presentato et al., 2020a), potentially suggesting a partial loss of flavonoids upon filming procedure. The detection of IR absorption bands accountable for -CH_2_ bending vibrations of aromatic compounds (ca. 1512 cm^-1^) for IPFs (Figure S1; Table 1) further indicated the occurrence of carotenoids and phenols within these films (Lóránd et al., 2002; Ricci et al., 2015; Rashid, Kait, & Murugesan, 2016), which were instead absent in CPF (Table S1).

Although a high similarity in vibrational modes was observed in the 1800-1480 cm^-1^ region for IP powders (Presentato et al., 2020a) and films (Table 1), the deconvolution of the latter revealed important differences, likely caused by the filming process itself. In this regard, IPFs lacked the large IR absorbance at ca. 1595 cm^-1^ attributable to aromatic skeleton vibrations (Heredia-Guerrero et al., 2014; Ricci et al., 2015; Rashid et al., 2016) that was detected for powdered formulations (Presentato et al., 2020a). This result may (*i*) corroborate the loss of volatile and bioactive substances during the filming of IPs or (*ii*) indicate an interaction of these compounds with pectin, Tween 60, and/or glycerol, which could cause a shift of IR bands that may eventually overlap with other vibrational modes. An additional variation between IPFs and powder formulations was the arising, in the former, of the asymmetric -COO^-^ stretching contribution of carboxylate groups within polygalacturonic acid (1605 and 1610 cm^-1^) (Figure 1; Table 1). A similar result was observed for CPF (Figure 1), suggesting that this modification may derive from the filming process. Indeed, upon dissolution of CP or IPs in water, -COOH moieties undergo dissociation, causing the arising of a negative charge distribution deriving from the large pool of -COO^-^ groups (Wellner, Kacurakova, Malovikova, Wilson, & Belton, 1998; Manrique & Lajolo, 2002), which are almost absent in the powders. These modifications are directly linked to the degree of esterification (DE) (*i.e.*, the fraction of -COOH groups esterified with methanol) of pectin polymers (Fidalgo et al., 2016; Wang et al., 2016; La Cava et al., 2018) here investigated, which is reported in **Table 2**. Indeed, the lower the DE of pectin, the higher is the number of -COOH groups that can dissociate in their anionic forms in water (Manrique & Lajolo, 2002).

**Table 2.**
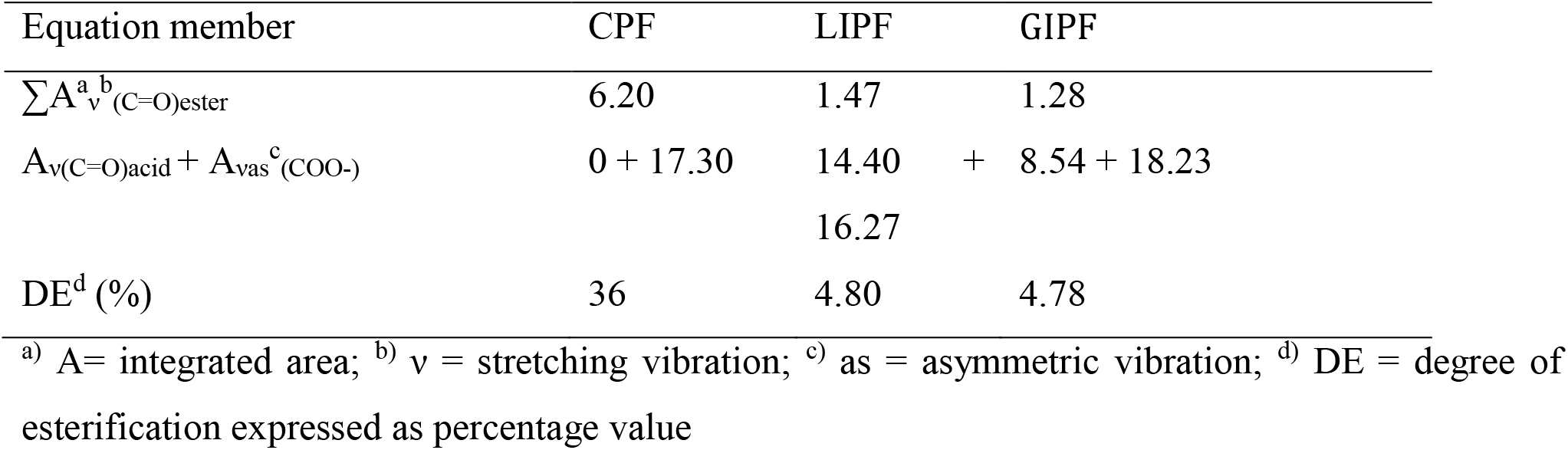
Degree of Esterification (DE) of CP, LIP, and GIP films.

Following the Fidalgo-Ilharco equation (section 2.3), the films contained low methoxy (LM) pectins, as their DE was lower than 50% (Table 2). This feature is of paramount importance for film generation, as LM-pectin can easily produce gels upon adding divalent cations (Ca^2+^ in this study) that can interact with a large amount of free -COO^-^ groups within a broad pH range (Wellner et al., 1998; Günter & Popeyko, 2016). Nevertheless, CPF featured a higher DE value (36%) than IPFs (ca. 4.80%), agreeing with our observations regarding the powder formulations (Presentato et al., 2020a). This difference links to the better performance of the HC used for IP extraction in preserving hydrophilic RG-I chains than acidic hydrolysis (for CP) (Kaya, Sousa, Crepeau, Sorensen, & Ralet, 2014; Wang et al., 2017a). Indeed, these pectin chains have a strong preference to form gels by addition of Ca^2+^ (Günter et al., 2016), as suggested by the IR band centered at ca. 1410-1420 cm^-1^, which can trace back to the symmetric stretching of pectin -COO^-^ groups interacting with Ca^2+^ (Wellner et al., 1998).

The so-called polysaccharide IR “fingerprint region” (1200-950 cm^-1^) showed intense and convoluted bands mainly related to the crystallinity and conformation of pectin polymers (Figure 1; Table S1). Besides, IR absorbance deriving from the presence of bioactive substances (*i.e*., flavonoids, terpenes, and terpenoids) was detected in IPFs’ ATR-FTIR spectra (Figure 1; Table S1). To better resolve the contributions within this region, spectral deconvolutions were performed (**Figure S2**; **Table 3**).

**Table 3.** Deconvolution of CPF and IPFs spectra by non-linear least squares fitting of the 1200-950 cm^-1^ region.

| CPF |  |  | LIPF |  |  | GIPF |  |  | Vibrational mode<sup>a</sup> |
| --- | --- | --- | --- | --- | --- | --- | --- | --- | --- |
| $\tilde{\nu} [\text{cm}^{-1}]^b$ | w<sup>c</sup> | A<sup>d</sup> | $\tilde{\nu} [\text{cm}^{-1}]$ | w | A | $\tilde{\nu} [\text{cm}^{-1}]$ | w | A | $\nu^e(\text{C-O-C})_{\text{glycosidic bond}};$<br>$\nu (\text{C-C})_{\text{pectin ring}}$ |
| 1142 | 31.7 | 7.97 | 1145 | 16.6 | 0.94 | 1144 | 17.7 | 1.23 | $\nu (\text{C-O}); \nu (\text{C-OH});$<br>$\nu (\text{C-O-C}); \nu (\text{C-C})$ |
| 1108 | 16.5 | 4.61 | 1104 | 27.1 | 11.9 | 1105 | 28.3 | 10.9 | $\nu (\text{C-O}); \nu (\text{C-OH});$<br>$\nu (\text{C-O-C}); \nu (\text{C-C})$ |
| 1095 | 16.4 | 5.29 | | | | | | | $\nu (\text{C-O}); \delta^f (\text{OH})$ |
| 1082 | 7.35 | 0.50 | | | | | | | $\nu (\text{C-O}); \rho^g (\text{CO});$<br>$\nu (\text{C-O-C}); \nu (\text{C-C});$<br>$\nu (\text{C-OH})$ |
| 1073 | 11.7 | 1.48 | 1071 | 30.6 | 10.5 | 1072 | 37.0 | 12.7 | $\nu (\text{C-O}); \rho (\text{CO}); \nu$<br>$(\text{C-O-C}); \rho (\text{CH}_3);$<br>$\nu (\text{C-OH}); \nu (\text{C-C})$ |
| 1048 | 75.8 | 26.2 | 1045 | 20.7 | 9.09 | 1045 | 20.1 | 4.06 | $\nu (\text{C-O}); \nu (\text{C-O-C});$<br>$\nu (\text{C-C}); \nu (\text{C-OH});$<br>$\rho (\text{CO});$<br>$\rho (\text{CH}_3)_{\text{flavonoids}}$ |
| | | | 1025 | 24.8 | 16.2 | 1024 | 49.6 | 13.4 | $\nu (\text{C-C}); \nu (\text{C-O})_{\text{pectin}}$ |
| 1014 | 15.2 | 9.09 | 1011 | 14.9 | 1.80 | | | | $\nu (\text{C-C}); \nu (\text{C-O})$ |
| 1006 | 8.48 | 0.33 | | | | | | | $\nu (\text{C-O});$<br>$\nu (\text{C-C})_{\text{glycosidic bond}};$<br>$\nu (\text{H}_6\text{C-C})_{\text{pyranosyl ring}}$ |
| | | | 995 | 21.4 | 7.43 | 992 | 16.6 | 2.81 | $\nu (\text{C=C})_{\text{trans}};$<br>$\gamma^h (=CH);$<br>$\rho (\text{CH}_3)_{\text{flavonoids}}$ |
| 971 | 42.8 | 6.83 | 973 | 23.9 | 5.71 | 973 | 25.1 | 5.72 | $\delta$ (C=O); $\delta$ (CCH);<br>$\delta$ (COH) |
| 958 | 19.1 | 1.35 | | | | | | | $\delta$ (C=O); $\delta$ (CCH);<br>$\delta$ (COH) |

<sup>a)</sup> references are enlisted in the Supplementary Material; <sup>b)</sup> $\tilde{\nu}$ [ $\text{cm}^{-1}$ ] = wavenumber; <sup>c)</sup> w = width; <sup>d)</sup> A = integrated area; <sup>e)</sup> $\nu$ = stretching vibration; <sup>f)</sup> $\delta$ = bending vibration; <sup>g)</sup> $\rho$ = rocking vibration; <sup>h)</sup> $\gamma$ = out of plane ring vibrations.

Spectral deconvolutions revealed a higher signal variability for CPF than IPFs likely to ascribe to the partial overlapping, in the latter, of polysaccharide contributions with those deriving from bioactive substances (Table 3; Table S1). IR absorption bands shared between all films referred to vibrational modes typical of the pyranose ring, glycosidic bond, alongside arabinose and galactose residues (Presentato et al., 2020a). A larger integrated area of IR contributions attributable to the pectin ring and its glycosidic bonds (ca. 1140 cm^-1^) was observed for CPF than IPFs (Table 3), suggesting the major contribution of the polysaccharide in the former (Presentato et al., 2020a). In line with our previous investigations (Presentato et al., 2020a), neutral sugars arabinose and galactose (ca. 1070 and 970 cm^-1^) were more represented for IPFs than CPF (Table 2), being the largest area calculated for GIPF, as galactose is among the most present neutral sugars found in grapefruits (Kaya et al., 2014). Besides, IPFs featured IR bands related to the presence of terpenoids (ca. 1105 cm^-1^), flavonoids (ca. 1025 and 970 cm^-1^), and terpenes (ca. 990 cm^-1^) (Schulz & Baranska, 2007; Presentato et al., 2020a; Nogueira et al., 2020; Shivangi et al., 2021) that were either absent or less represented in CPF (Table 3), where these signals derived from the overlapping vibration of the pectin polymer. Vibrational modes of glycerol and Tween 60 were more represented in IPFs than CPF (Table 2; Table S1). This phenomenon can depend on the interaction of bioactive substances within IPFs with the pectic material that can diminish the pectin moieties available to interact with the plasticizer and surfactant. Comparing these spectral deconvolutions with those obtained for pectin powders (Presentato et al., 2020a) highlighted the occurrence of critical differences between ATR-FTIR spectra, hence the physical-chemical nature of these formulations. CP and IP filming led to the arising of an IR band centered at ca. 1105 cm^-1^ (Table 3; Table S1), which can relate to the presence of glycerol, bioactive substances in IPFs, and a modification of the pectin structure itself. The latter hypothesis was further suggested by the smaller integrated areas obtained for IR signals typical of C-C and C-O stretching vibrations (ca. 1140 and 1015 cm^-1^) deriving from the pectin ring, glycoside bond, and polygalacturonic acid residues in the case of films (Table 2) than powder formulations (Presentato et al., 2020a). This result might derive from the crosslinking action exerted by Ca^2+^, which can interact with oxygen atoms present in (*i*) - COO^-^ groups (O_6_), (*ii*) the pectin ring (O_5_), (*iii*) the glycosidic bond (O_4_-O_1_’), and (*iv*) -OH groups of adjacent galacturonic residues (O_2_’) (Wellner et al., 1998), causing important modifications within the 1150-1000 cm^-1^ region of ATR-FTIR spectra (Figure 1 and Table 3; Figure S2 and Table S1).

IR absorption bands detected for pectin films between 950 and 650 cm^-1^ were attributable to pectin vibrations and, in the case of IPFs, bioactive substances (Table S1). Particularly, these contributions were largely similar to those of powder formulations (Presentato et al., 2020a), exception for the arising, in CPF, of an IR signal referring to -COO^-^ bending vibration (ca. 770 cm^-1^; Table S1), which is attributed to the partial esterification of polygalacturonic acid residues (Bichara et al., 2016).

#### 3.1.2 ^13^C CPMAS NMR spectroscopy

^13^C CPMAS NMR spectra of CPF and IPFs showed signals typical of the pectin polymer, whose intensity was higher for the former than IPFs (**Figure 2; Table S2**), likely deriving from the higher amount of this polymer in the CPF. On the opposite, the absence of resonance peaks attributable to bioactive substances may derive from their low concentrations within IPFs, which impaired their identification (Liu et al., 2020).

**Figure 2.**
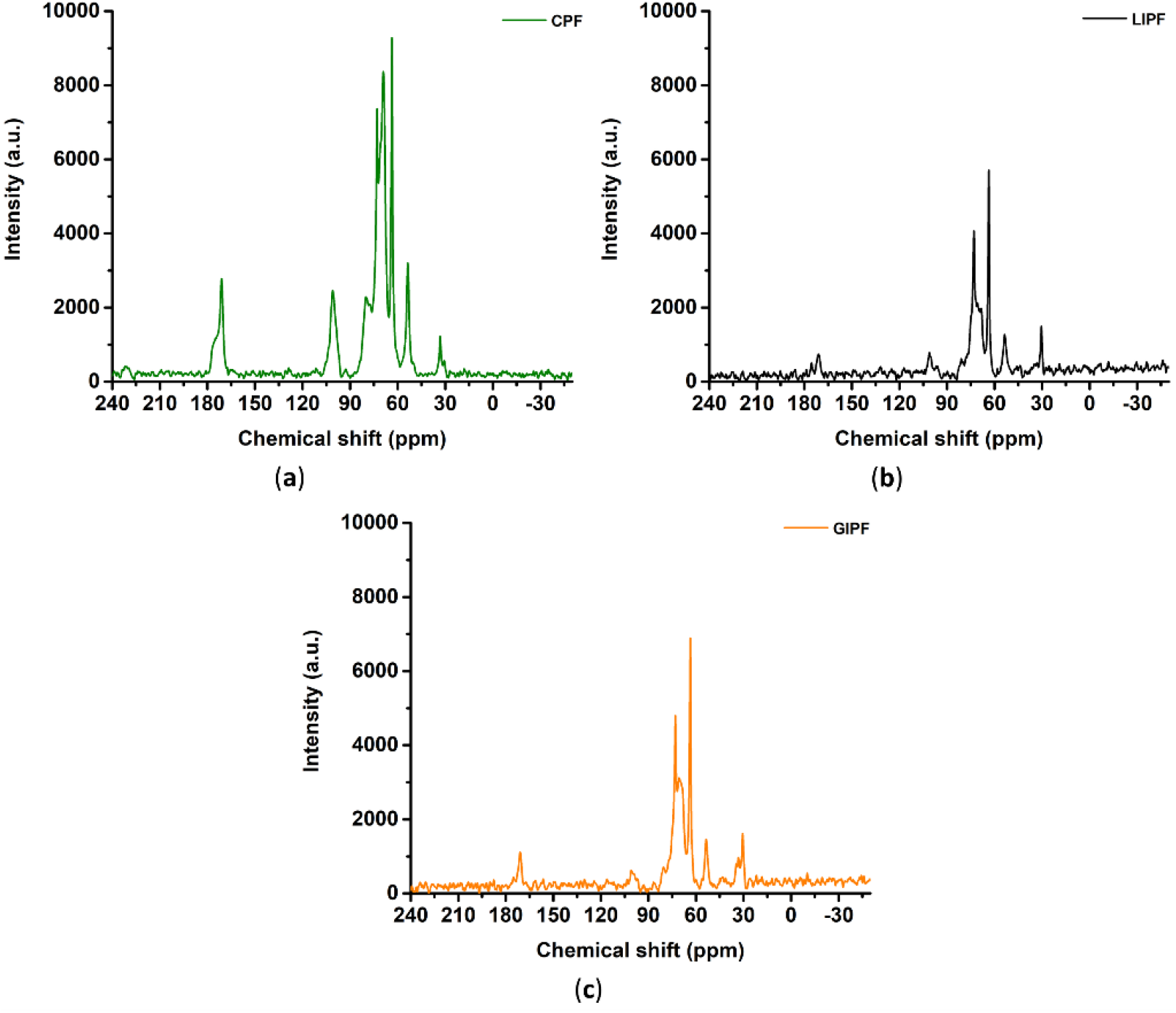
^13^C CPMAS NMR spectra collected for CPF (a), LIPF (b), and GIPF (c).

Chemical shifts at 177-170 ppm, 101-100 ppm, 81-77 ppm, and 73-69 ppm were attributed to C6 of galacturonic acid units, C1 and C4 in the glycosidic bond of pectin, and C2, C3, C5 of the pyranoid ring (Table S2) (Synytsya, Copikova, & Brus, 2003), respectively. All films showed additional signals related to the pectin polymer at ca. 53.6 ppm (Figure 2; Table S2), which indicated the methyl esterification of C6 (Synytsya et al., 2003), and an intense signal (ca. 63.6 ppm) deriving from glycerol (Soltys et al., 2019) (Figure 2; Table S2). In this regard, although one of the typical C peaks of glycerol is centered at 66.9 ppm (Breitmaier & Voelter, 1974), its interaction with pectin, alongside Tween 60 and, in the case of IPFs, bioactive substances, may have caused a downshift of this signal. Indeed, both the physical and chemical environment of glycerol can undergo substantial modifications due to the surrounding and interacting substances. The second C signal generally attributed to glycerol molecules (ca. 77 ppm) likely overlapped with chemical shifts of the pyranoid ring (Soltys et al., 2019), making its identification unlikely. CPF and IPFs also displayed C peaks referring to (-(CH_2_)_n_) presence (33-30 ppm; Figure 2; Table S2), most likely deriving from Tween 60. Nevertheless, this surfactant interacted differently with CPF and IPFs, as highlighted by the change in the 33ppm/30.5ppm intensity ratio (Figure 2). The higher intensity of 30.5 ppm signal observed for IPFs might derive from the occurrence of physical-chemical interactions between the surfactant and hydrophobic moieties of bioactive substances contained in IPs, which were absent in the case of commercial pectin.

An important difference between IPFs and CPF was the detection, in the former, of weak chemical shifts in the 43-45 ppm range. This signal can link to fully substituted quaternary carbons typical of terpenoid mixtures, such as essential oils (Conte et al., 2010), further confirming the presence of bioactive substances in IPFs. Similarly, C1 resonance differed between CPF, which displayed a ^13^C NMR singlet centered at ca. 101.2 ppm, and IPFs, for which a doublet (ca. 101 and 99 ppm) appeared (Figure 2; Table S2). The latter suggested more ordered chains in pectin of IPFs than CPF, most likely consequently to both glycerol addition (Soltys et al., 2019) and bioactive substances interacting with the polymer. The finding agrees with ATR-FTIR spectroscopy results (Figure 1; Table S1), which highlighted signals of glycerol. Furthermore, typical non-interacting glycerol vibrational modes were detected for IPFs (Figure 1; Table S1), thus suggesting a competitive mechanism of interaction among this plasticizer and bioactive substances. Signals in the 180-170 ppm region attributed to carboxyl C6 of galacturonic units (Gal) varied between films, indicating the presence of diverse substitutions at this position (Synytsya et al., 2003), which supports the type III interaction mechanism among pectin and glycerol involving the carboxylic group (Vityazev et al., 2020). For instance, CPF featured a resonance peak at ca. 177 ppm that belonged to -COO^-^ moieties of pectin (Synytsya et al., 2003), which was absent for IPFs, while carbonyl carbons of acetyl groups (OCOCH_3_; ca. 171.3 ppm) (Synytsya et al., 2003) were detected for all pectin-based films (Figure 2; Table S2). Acetylation results in marked variations of pyranoid ring carbons, whose resonance is registered between 76 and 66 ppm (Synytsya et al., 2003). Thus, spectral deconvolutions were performed in this region to study better the films’ nature (**Table 4; Figure S3**).

**Table 4.**
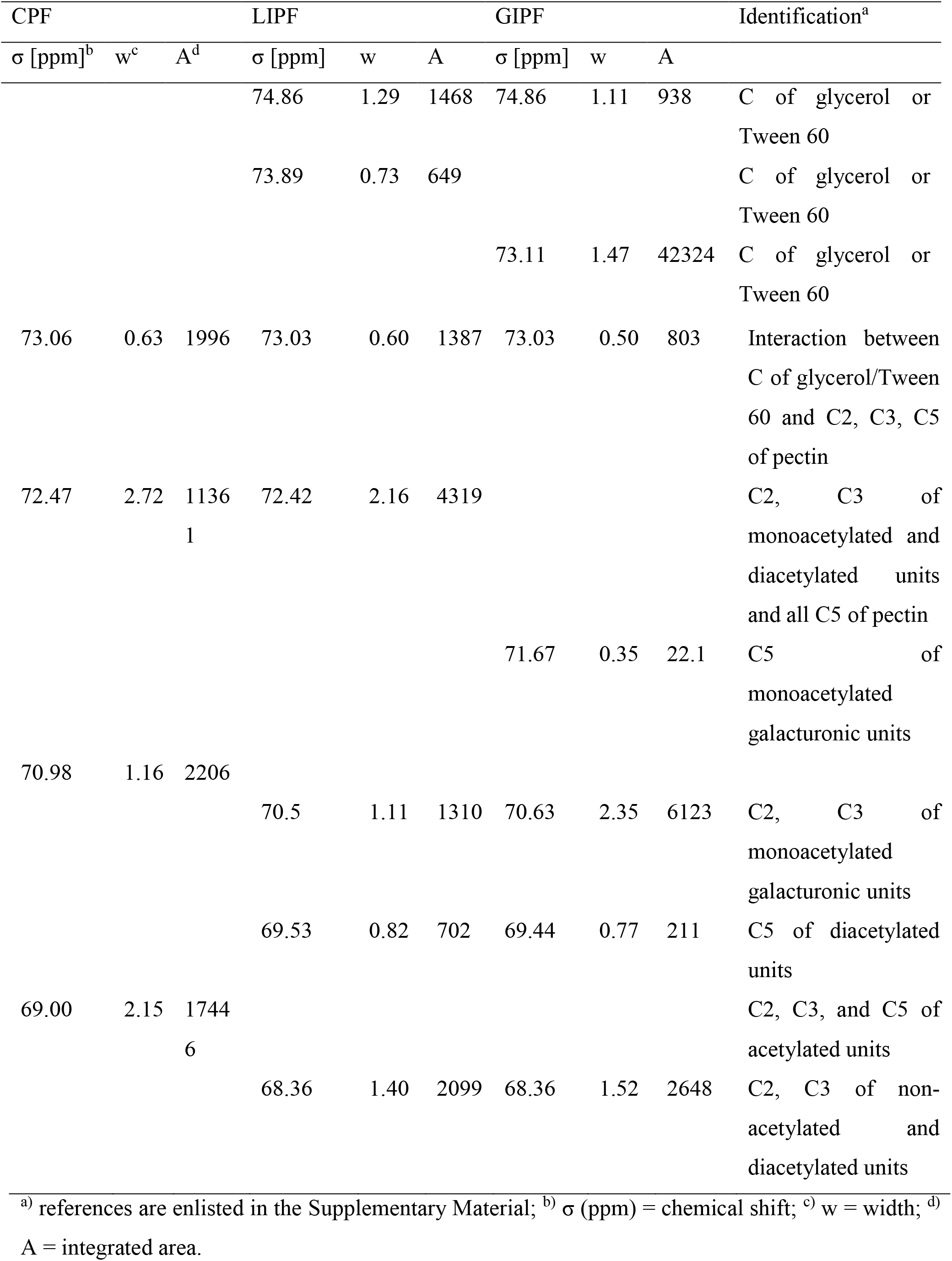
Spectral deconvolution by non-linear least squares fitting of the 76-66 ppm region of ^13^C CPMAS NMR spectra.

The greater complexity in composition of IPFs than CPF reflected the higher variability of resonance contributions observed for the former through ^13^C CP MAS NMR deconvolution (Table 3; Figure S3). Indeed, bioactive substances, glycerol, and Tween 60 may have caused the arising of interactions strongly impacting the polymeric environment within IPFs. When aromatic compounds, such as phenols and polyphenols, interact with polysaccharides, variations in the electron cloud density of both the aromatic ring and the carbohydrate moiety occur, which are detectable through variations in their resonance peaks (Liu et al., 2020). Specifically, the more complex the phenolic molecule is, the greater is the chemical shift variation caused by its interaction with polysaccharides (Liu et al., 2020). Considering that IPs contained a mixture of aromatic and phenolic compounds (Presentato et al., 2020a; Scurria et al., 2021a; Scurria et al., 2021b), this phenomenon should be emphasized for IPFs than systems featuring only one type of phenolic molecule. Moreover, the same bioactive substances, alongside pectin, may interact with glycerol and Tween 60, causing, for instance, a shift or a modification of their resonance peaks typically observed between 76 and 66 ppm, further explaining the diversity of IPF and CPF deconvolution results (Table 3). IPFs also showed diverse C2, C3, and C5 NMR contributions than CPF (Table 3; Figure S3), which suggested a higher acetylation in the former. Indeed, a larger integrated area was detected in the 72-70 ppm range for IPFs, while CPF displayed more prominent contributions in the 70-69 ppm region, which indicated a higher DE (Synytsya et al., 2003) corroborating our ATR-FTIR results. Although CPF featured resonance peaks attributable to acetylated Gal (72.47 and 69.0 ppm), these contributions were more represented for IPFs (Table 3). Particularly, spectral deconvolution of IPFs displayed NMR signals centered at ca. 70.6 and 69.5 ppm, which are typical of C2 and C3 of monoacetylated Gal units and C5 of diacetylated Gal, respectively (Synytsya et al., 2003), while these contributions were absent in CPF (Table 3). Additional resonance peaks attributable to C2 and C3 diacetylated pectin units were identified for IPFs at 68.36 ppm (Table 3) (Synytsya et al., 2003). Yet only GIPF displayed a weak contribution linked to C5 of monoacetylated ones at 71.6 ppm (Table 3) (Synytsya et al., 2003). From an applicative perspective, the higher acetylation of IPFs than CPF can be advantageous, as acetylated pectins feature a higher interfacial and surface tension, which, in turn, makes them good emulsifying agents (Yang, Mu, & Ma, 2018).

#### 3.1.3 XRD analysis

The amorphous or crystalline nature of CP and IPs was determined through XRD, focusing on both powder and film formulations to investigate potential modifications deriving from the filming procedure (**Figure 3**).

**Figure 3.**
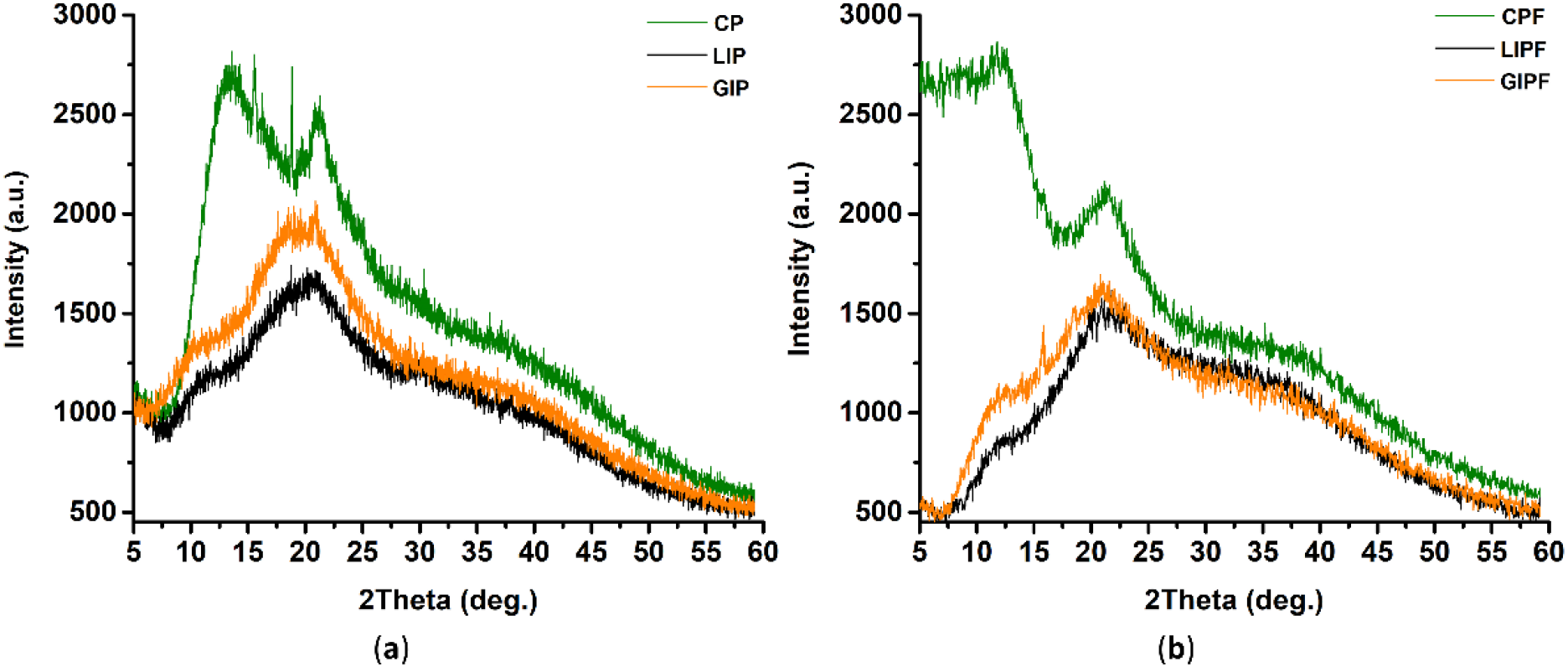
XRD patterns of pectin-based powder formulations (a) and films (b).

The diffraction patterns of pectin powders indicated a partial crystalline structure of the three samples (Figure 3 a), as CP, LIP, and GIP featured a degree of crystallinity (DC) of 37, 24, and 22%, respectively. In line with previous studies (Meneguin, Cury, & Evangelista, 2014; Wang et al., 2016; Chaichi, Hashemi, Badii, & Mohammadi, 2017), CP showed diffraction peaks centered at 13.65 and 21.26° (Figure 3 a) that could relate to second order reflections of helical structure proposed for pectin by Walkinshaw and Arnott (1981a). IPs’ bioactive substances appeared to modify the crystalline structure of pectin, as highlighted by the shift observed for both first and second peaks (Figure 3 a). Moreover, two additional contributions appeared at 16.11 and 18.89°, yet the latter had a lower intensity in LIP than GIP (Figure 3 a). In this regard, polyphenols and phenolic acids can generate hydrogen bonds with pectin residues, reducing the pectin intermolecular attraction (Nisar et al., 2019). This event, in turn, can cause a modification (or even the disruption) of the internal crystal structure of the polymer itself, diminishing its crystallinity (Peng, Wang, Shi, Chen, & Zhang, 2020).

Upon filming, XRD patterns underwent some modifications (Figure 3 b). The peak of CP powder centered at 16.11° shifted to 12.41° in CPF (Figure 3 b), thus indicating a variation in the lateral spacing among pectin chains (Walkinshaw & Arnott, 1981a), likely due to the addition of Ca^2+^ and Tween 60. Indeed, bridges formed between pairs of pectin chains can result in the generation of different junction zones in the polymer (Walkinshaw & Arnott, 1981b), while Tween 60 was previously reported to cause complex variations in the crystalline conformation of the polysaccharide-based film (Peng et al., 2020). IPF diffraction patterns indicated structural variations comparable to CPF (Figure 3 b). Nevertheless, the DC increased for IPFs (36 and 32% for LIPF and GIPF, respectively), confirming the ability of glycerol to act as a plasticizer likely through a redistribution of the crystal type within the polymeric matrix (Ziani, Oses, Coma, & Maté, 2008). This event can promote a more ordered structure, corroborating the modification of the C1 resonance peak observed for IPFs (Figure 2). The DC of CPF did not show significant variation (34%), being in line with the indication of a disordered phase obtained by analyzing the C1 NMR signal (Figure 2). In this regard, although the same amount of glycerol was used for all films, the addition of Tween 60 to CP may have led to a strong modification of the polymer crystal structure. In IPFs, bioactive substances limited this phenomenon, as they can preferentially interact with this surfactant through hydrogen bonds or their hydrophobic moieties.

### 3.2. Release of bioactive substances from IPFs

Both IPFs showed the ability to act as carrier and controlled-release systems for bioactive substances absorbing light in the UV region over time (2-72h), yet some differences arose (**Figure 4).**

**Figure 4.**
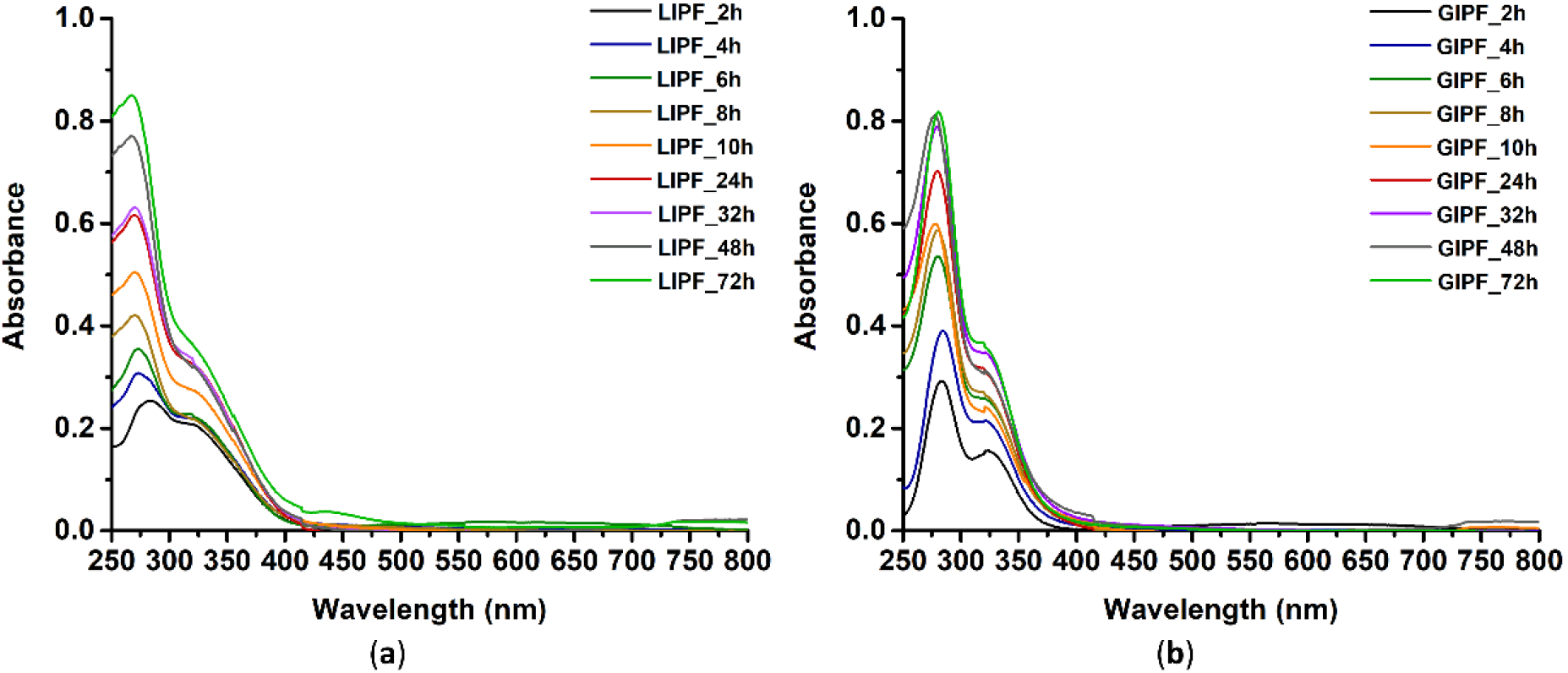
UV-Visible spectra of LIPF (a) and GIPF (b) incubated in TSB medium from 2 to 72h.

Absorbance maxima for LIPF were detected in the 267-283 nm region, while GIPF featured maximum centered at ca. 283 nm over the entire timeframe (Figure 4). Specifically, the main absorbance peak of LIPF blue shifted from 281 (2h incubation) to 267 nm (72h of incubation; Figure 4), suggesting that this film may be releasing different bioactive substances over time. Both IPFs displayed an absorbance shoulder at ca. 323 nm, yet it was more resolved in LIPF (Figure 4). Besides, based on the measured absorbance intensities, GIPF seemed to release bioactive molecules faster than LIPF, reaching a plateau at 32h of incubation (Figure 4). On the opposite, the latter appeared to mediate a slow yet more controlled release, as indicated by the constant increase in absorbance value up to 72h of incubation (Figure 4).

Absorbance contributions were analyzed in detail by deconvolution of UV-Visible spectra focusing on the 250-500 nm region (**Figures S4** and **S5**; **Table S3**). Most of these contributions were attributable to phenolic compounds within IPFs (Presentato et al., 2020a). This observation may indicate the preferential release of polyphenols, such as phenolic acids and flavonoids, in an aqueous environment from the pectin polymer, likely due to their higher solubility in water than volatile compounds such as terpenes and terpenoids. Furthermore, polyphenols tend to develop non-covalent interactions (*i.e.*, hydrogen bonds or hydrophobic interactions) with polysaccharides, favoring the release of these bioactive substances (Zhu, 2021). Similar interactions generally occur between polyphenols and plasticizers or surfactants, partially contributing to their release in an aqueous environment.

In line with our previous reports (Scurria et al., 2021a; Scurria et al., 2021b), LIPF showed more variability in absorbance peaks than GIPF, indicating the presence of a higher number of bioactive substances in the former (Figures S4 and S5; Table S3). Based on compositional analyses previously performed on IPs (Scurria et al., 2021a; Scurria et al., 2021b), a putative identification of absorbance contributions was performed. The absorbance at 270-275 nm was ascribed to gallic acid (Fink & Strong, 1982; Luna et al., 2016), kaempferol (Tian, Liu, Tian, Hu, & Chen, 2004), and kaempferol-7-O-glucuronide (Carazzone, Mascherpa, Gazzani, & Papetti, 2013), while naringin (Roy, Tripathy, Chatterjee, & Dasgupta, 2010), naringenin (Yousuf & Muthu Vijayan, 2013), hesperidin (Srilatha, Nasare, Nagasandhya, Prasad, Diwan, 2013), and eriocitrin (Lauro et al., 2017) were responsible for the 280-288 nm contributions. These phenolic compounds also featured absorbance shoulders in the near UV region, allowing their identification in the spectral deconvolutions (Table S3). Indeed, (*i*) kaempferol and its glucuronide form display a typical shoulder in the 360-370 nm range (Tian et al., 2004; Carazzone et al., 2013), (*ii*) naringin, naringenin, and eriocitrin at ca. 320-325 nm (Roy et al., 2010; Yosouf & Muthu Vijayan, 2013; Lauro et al., 2017), and (*iii*) hesperidin showed a shoulder centered at ca. 350 nm (Srilatha et al., 2013). Finally, the additional absorbance peaks observed for LIPF between 304 and 318 nm were attributed to *p*-coumaric acid (Putschogl, Zirak, & Penzkofer, 2008) and the monoterpenoid safranal (Kanakis, Tarantilis, Pappas, Tajmir-Riahi, & Polissiu, 2009). Specifically, GIPF contained, for the most part, naringin and naringenin, while no eriocitrin was detected (Scurria et al., 2021b). Thus, the large and principal absorbance observed for GIPF at 280-288 and 320-325 nm (Figure 4; Figure S5; Table S3) was likely due to the high amount of naringin and naringenin present in GIP powder. On the opposite, the latter flavonoids were either absent or present in low amounts in LIP (Scurria et al., 2021b), suggesting that the same absorbance peaks (Figure 4; Figure S4; Table S3) traced back to eriocitrin in LIPF. Similarly, kaempferol and its glucuronide form, gallic acid, *p*-coumaric acid, and hesperidin, were present/more represented in LIP than in GIP, confirming the larger contributions in absorbance of these bioactive substances for LIPF (Figure 4; Figures S4 and S5; Table S3).

#### 3.2.1 Kinetics profile of bioactive substances released from IPFs

The first-order, second-order, and Ritger-Peppas models yielded good data fits for absorbance contributions of both IPFs (**Figure 5** and **Table S4**), while the remaining release models resulted in non-significant fitting of UV-visible data.

**Figure 5.**
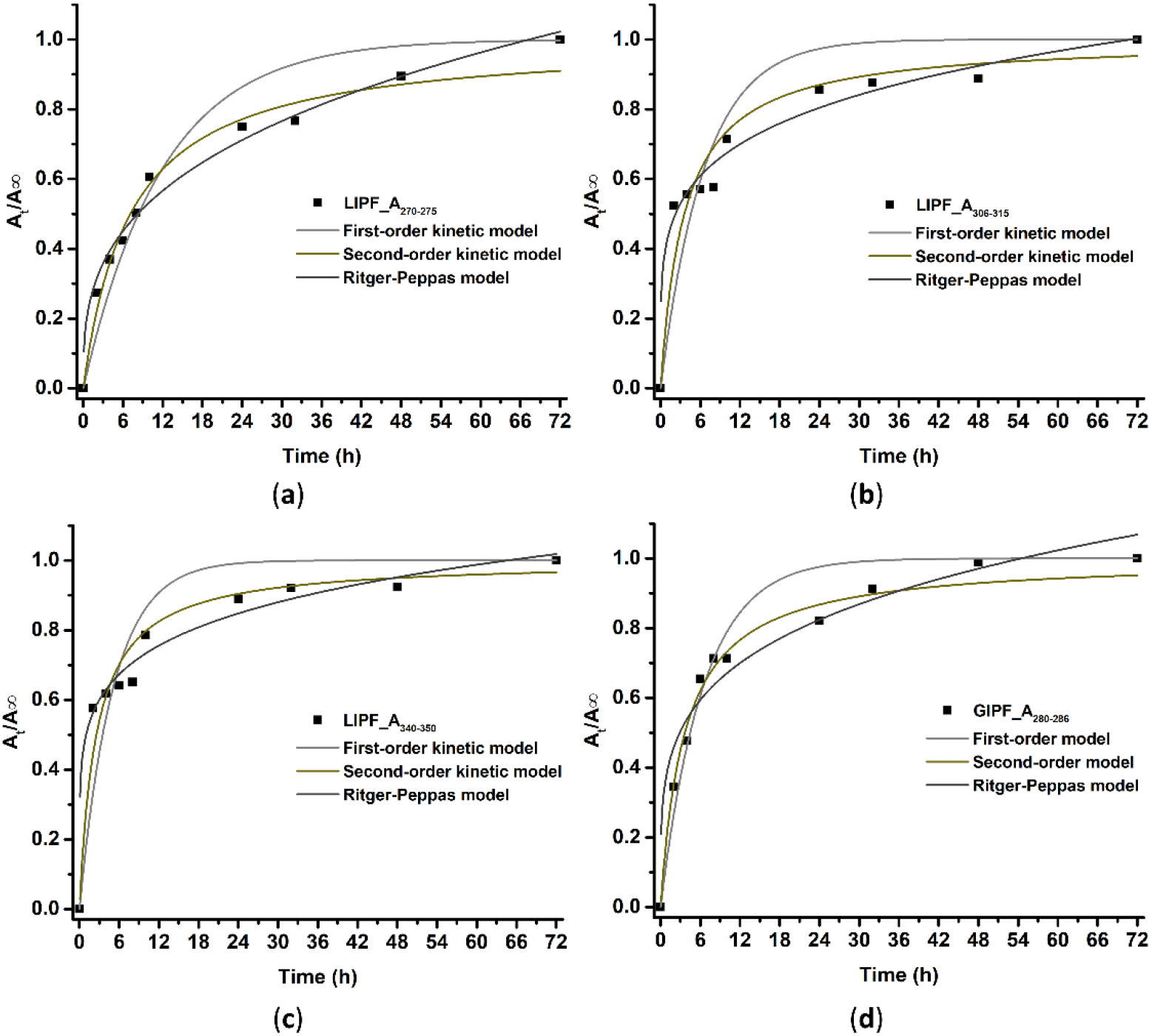
Release profiles obtained through first-order, second-order, and Ritger-Peppas models indicating the release of bioactive substances obtained for IPFs incubated in TSB medium from 2 to 72h. Kinetic profiles were derived from LIPF absorbance contributions in the 270-275 nm (a), 306-315 nm (b), and 340-350 nm (c) range, and those of GIPF in the 280-285 nm region.

The highest R^2^ values were obtained through the Ritger-Peppas model for LIPF contributions, while GIPF absorbance better fitted with the second-order kinetic profile (R^2^ = 0.9680) than the former (R^2^ = 0.9145) (Table S4). These results indicated that diffusion processes (second-order kinetic and Ritger-Peppas models) controlled the release of bioactive substances from IPFs, whose rate was proportional to the content of substances within the films (first-order model) (Yong et al., 2020). Following the release profiles described by the Ritger-Peppas model, a release exponent n = 0.5 corresponds to a Fickian diffusion mechanism, while when n > 0.5 non-Fickian phenomena relying on swelling or relaxation of the polymer chains occur (Ritger & Peppas, 1987). In IPFs, quasi-Fickian diffusion mechanisms appeared to mediate bioactive substance release, as n < 0.5 was obtained for all absorbance contributions (Liu et al., 2018; Jridi et al., 2019; Wu et al., 2020) (Table S4).

IFPs showed similar bioactive substance release profiles, where two phases were identified. At first (0-8h of incubation), a fast release of these compounds was detected for both IPFs, likely deriving from bioactive substances close to film surfaces (Liu et al., 2018; Gao et al., 2019; Wu et al., 2020). Subsequently (up to 72h), a sustained, controlled, and slower release was detected (Figure 5), as bioactive compounds immobilized by the pectin polymer into the inner core of IPFs may need more time to diffuse into the water environment (Liu et al., 2018; Gao et al., 2019; Wu et al., 2020). Moreover, although an initial “burst phase” was detected (Figure 5), this phenomenon spread over 8h of IPFs’ incubation in TSB medium, indicating a good retention of bioactive substances and their gradual release by these films. In line with this hypothesis, no constant (*plateau*) phase of substance release was observed for LIPF (Figure 5), suggesting that it could more finely control this phenomenon over time. The natural potentiality of pectin as a delivery polymer may also have been boosted by the addition of glycerol and Tween 60. Indeed, these compounds were previously described to slow down the release of polyphenolic extracts due to their effect on the crystal structure and solubility of polysaccharide film (Huaman-Castilla et al., 2020; Peng et al., 2020).

### 3.3 Antimicrobial activity of pectin-based films

Adherent growing cells challenged with IPFs showed inhibition halos qualitatively indicating how these films can elicit an antimicrobial activity against *P. aeruginosa* strains, whereas virtually no effect was detected against *K. pneumoniae* growth (**Figure S6**). IPFs more efficaciously inhibited *P. aeruginosa* ATCC 10145 than the other bacterial strains (Figure S6) likely due to their higher intrinsic tolerance and/or resistance towards antimicrobials. Bigger inhibition halos were noticed upon *P. aeruginosa* exposure to LIPF than GIPF (Figure S6), suggesting that the former could be more prone to exert an antimicrobial effect. IPFs’ relatively low efficacy in preventing adherent cell growth may ascribe to the low diffusion ability of bioactive substances on solid (or hydrogel) medium. Indeed, the release of these compounds from polysaccharide films mostly relies on swelling induced processes (Nogueira et al., 2020). When films are placed in a compatible medium (in our case water), the latter enters the polysaccharide matrix, causing the film to swell. In turn, the diffusion coefficient of bioactive substances within the films increases and the latter are more prone to diffuse out (Nogueira et al., 2020; Zhu, 2021). This phenomenon can be enhanced by the tendency of bioactive compounds such as polyphenols to form non-covalent (less strong) bonds with the polysaccharidic matrix (Zhu, 2021).

In line with our suggestion of a swelling release of bioactive substances from IPFs, the latter more efficiently prevented bacterial growth in the form of planktonic cells incubated in TSB medium (**Figure 6**).

**Figure 6.**
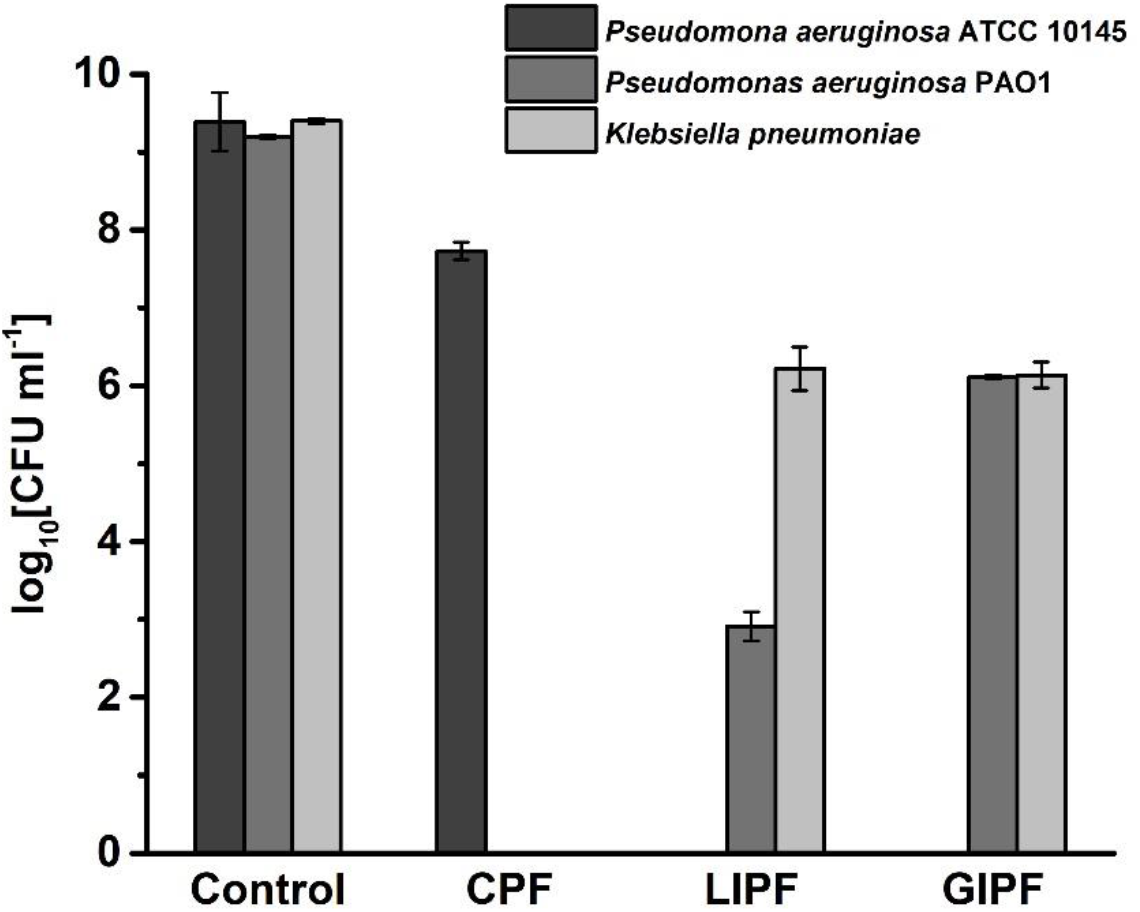
Antimicrobial activity of LIPF and GIPF against planktonically growing *Pseudomonas aeruginosa* ATCC 10145, *Pseudomonas aeruginosa* PAO1, and *Klebsiella pneumoniae* cells. Given the high resiliency of the two latter strains, the commercial *Citrus* film has been solely tested against the less aggressive *Pseudomonas aeruginosa* ATCC 10145.

CPF minimally inhibited *P. aeruginosa* ATCC 10145 (Figure 6), highlighting a small decrease in the number of CFU ml^-1^ (from 1·10^9^ - unchallenged cells - to 4.5·10^7^ - CPF exposed cells). Thus, given the intrinsic resistance of *P. aeruginosa* PAO1 and the clinical isolate to antimicrobials, the efficacy CPF was not assessed towards these bacterial strains. The minimal antimicrobial activity of CPF can derive from the oxidation of carbohydrate units within the pectin polymer by bacterial cells (Piacenza et al., 2018; Presentato et al., 2020a). This phenomenon may, in turn, trigger a series of cascade events (*i.e.*, turning-off of tricarboxylic acid cycle, intracellular accumulation of pyruvate, secretion and subsequent uptake of acetic acid, stimulation of the murein hydrolase activity) (Rice & Bayles, 2008) that eventually could compromise the cell viability. Both IPFs showed a higher ability in inhibiting bacterial cell growth than CPF (Figure 6), which traces back to the presence of bioactive substances well-known for their antimicrobial potential (Nazzaro, Fratianni, De Martino, Coppola, & De Feo, 2013). Specifically, IPFs acted as biocide against *P. aeruginosa* ATCC 10145, while the two other strains tolerated better the challenge represented by these film formulations (Figure 6). Nevertheless, the number of *K. pneumoniae* colonies decreased at least three orders of magnitude (> 99.9% of killing), in terms of logarithmic units, when challenged with LIPF (from 2.5·10^9^ to 1.83·10^6^ CFU ml^-1^) or GIPF (from 2.5·10^9^ to 1.43·10^6^ CFU ml^-1^) (Figure 6). A similar result was observed upon exposure of *P. aeruginosa* PAO1 cells to GIPF (from 1.60·10^9^ to 1.30·10^6^ CFU ml^-1^), yet the antimicrobial efficacy of LIPF was enhanced against this strain, as only an average of 8.5·10^2^ CFU ml^-1^ survived this challenge (Figure 6). A direct comparison between these results and those reported in literature is challenging due to the diversity in the bacterial strains used, the applied screening methods, solvents utilized for the antimicrobial testing, alongside concentration ranges and sources of bioactive substances explored (Adamczak, Ozarowski, & Karpinski, 2020; Zhu, 2021).

Based on our previous reports (Scurria et al., 2021a; Scurria et al., 2021b), the available literature, and the results here presented, some hypotheses regarding the mechanism of antimicrobial action of IPFs were formulated. Overall, flavonols, such as kaempferol and its glucuronide forms, are described for (*i*) their potent antimicrobial activity towards both Gram-negative and -positive bacterial strains, (*ii*) their synergism with antibiotics, and (*iii*) their ability to suppress several microbial virulence factors (Daglia, 2012). On the opposite, flavanones, such as naringenin, naringin, and hesperidin, more efficaciously inhibit Gram-positive growth, as these substances can more easily enter cells with a single-layered and low lipid-containing cell wall, damaging the membrane permeability, intracellular enzymes, and interfering with DNA synthesis (Celiz, Daz, & Audisio, 2011; Iranshahi, Rezaee, Parhiz, Roohbakhsh, & Soltani, 2015; Wang, Wang, Zeng, Xu, & Brennan, 2017b; Farhadi, Khameneh, Iranshahi, & Iranshahy, 2019). Hence, since GIPF mostly features naringenin and naringin (Scurria et al., 2021b), these compounds can only partially exert an antimicrobial effect towards Gram-negative bacterial strains. Yet, the boosted antimicrobial efficacy of LIPF can link to the presence of kaempferol and kaempferol-7-O-glucuronide (Scurria et al., 2021b) that were released over time (**Figure 5 a**). Indeed, kaempferol and its derivatives can disrupt the Gram-negative membrane by interacting with both the polar head and alkyl chains of phospholipids through their -OH and hydrophobic moieties, respectively (He, Wu, Pan, & Xu, 2014). Once inside the cell, these flavanols can exert their antimicrobial activity at different levels (Daglia, 2012) targeting key enzymes for the bacterial survival such as DnaB helicase of *K. pneumoniae* (Chen & Huang, 2011), dihydropyrimidinase of *P. aeruginosa* (Huang, 2015), and β-ketoacyl carrier protein synthase I and III of *Escherichia coli* (Lee, Lee, Jeong, & Kim, 2011). Kaempferol was also proven to generate Reactive Oxygen Species (ROS) through the interaction of its phenoxyl radical with oxygen or the reduction of iron and copper ions (del Valle et al., 2016). Besides flavonols, LIP featured a higher amount of phenolic acids (*i.e., p*-coumaric acid and gallic acid) and the monoterpenoid safranal than GIP (Scurria et al., 2021a; Scurria et al., 2021b), which are released the most over time by LIPF (**Figure 5a**). Phenolic acids have a higher affinity for Gram-negative bacteria than -positive ones due to the larger lipid content of the former. This feature can allow the integration of phenolic acids into the Gram-negative membrane, eventually leading to its disruption (Borges, Ferreira, Saavedra, & Simoes, 2013). In this regard, both *p*-coumaric acid and gallic acid can cause irreversible changes in cell membrane properties such as charge, permeability, and physical-chemical features (Lou et al., 2012; Borges et al., 2013). For instance, gallic acid can mediate (*i*) hydrophobicity variations, (*ii*) the decrease of membrane negative surface charge, (*iii*) the occurrence of local rupture, and (*iv*) the formation of pores in cell membranes, resulting in a substantial leakage of intracellular molecules in the extracellular environment (Borges et al., 2013). These phenolic acids can also act as permeabilizers increasing the permeability of both the outer and plasma membranes of *Salmonella, E. coli*, and *P. aeruginosa* spp., causing a drastic loss of barrier function (Simoes, Bennet, & Rosa, 2009; Lou et al., 2012; Abreu, McBain, & Simoes, 2012; Borges et al., 2013). The permeabilizer activity of gallic acid seems to link with its partial hydrophobicity and its ability to chelate divalent cations from the outer membrane (Abreu et al., 2012). Additional modes of action of gallic acid include (*i*) the inhibition of efflux pumps normally responsible for antimicrobial resistance mechanisms (Simoes et al., 2009; Abreu et al., 2012), (*ii*) the capability of acting as an electrophilic product for Gram-negative strains, affecting their electron acceptor ability (Borges et al., 2013), and (*iii*) the acidification of the cytoplasm that can result in protein denaturation (Borges et al., 2013). On the other hand, when *p*-coumaric acid enters bacterial cells, it can bind to DNA phosphate groups and intercalate the groove in the DNA double helix, affecting cellular replication and gene transcription (Lou et al., 2012). In the case of safranal, studies showed its antimicrobial efficacy against *Salmonella* and *E. coli* spp. (Pintado et al., 2011; Liu et al., 2017), yet its mode of action still needs to be elucidated. As a monoterpenoid, safranal may partition into the bacterial cell wall, causing, for instance, its fluidification or even disruption (Nazzaro et al., 2013). Moreover, Liu and colleagues (2017) reported on the ability of safranal to completely inhibit the ATP synthase in *E. coli*, which resulted in bacterial cell death. Thus, given the complexity of IPFs, their antimicrobial activity can derive from the ensemble of bioactive substances released over time, which, altogether, can synergistically act to inhibit bacterial growth. This hypothesis is particularly relevant for LIPF, as it showed higher variability in composition than GIPF (Scurria et al., 2021a; Scurria et al., 2021b), a greater antimicrobial efficacy, alongside a larger amount and release of flavonols, monoterpenoids, and phenolic acids known for their activity against Gram-negative strains.

## 4. Conclusions

LIP and GIP recovered from agri-food waste through HC constituted good candidates to produce bio- and eco-compatible films using glycerol, Tween 60, and aqueous CaCl_2_ as a plasticizer, surfactant, and cross-linking agent, respectively. A comparison between IPFs and CPF highlighted the uniqueness of the former. Indeed, the preservation of hydrophilic RG-I regions of IPs guaranteed a preferential interaction of polymer chains with divalent cations, forming stable gels. IPs also seemed to have a higher interfacial and surface tension than CP, which can favor the generation of stable emulsions.

A crucial added value of IPFs is the presence of bioactive substances, such as flavonoids, phenolic acids, terpenes, and terpenoids. These compounds interacted with pectin and glycerol through intermolecular hydrogen bonds, while hydrophobic moieties were likely involved in the case of Tween 60. These interactions, in turn, caused the generation of more ordered chains in the pectin of IPFs than CPF and a modification of the polymer crystal structure. Besides, IPFs acted as slow- and controlled-release systems of bioactive substances up to 72h through the occurrence of quasi-Fickian diffusion processes, being LIPF more efficient than GIPF. Specifically, phenolic acids, flavonols, and the monoterpenoid safranal were likely released from LIPF, while GIPF contained mostly flavanones. This intrinsic difference in composition and release reflected on the diverse antimicrobial potential of LIPF and GIPF against Gram-negative pathogens, which was null for CPF. Both IPFs acted as biocides against the pathogen indicator strain *P. aeruginosa* ATCC 10145, yet LIPF was more effective in inhibiting *P. aeruginosa* PAO1 growth. Moreover, IPFs substantially prevented the proliferation of the multidrug-resistant isolate *K. pneumoniae*. IPFs’ antimicrobial activity can trace back to the occurrence of diverse bioactive substances, which can synergistically limit pathogen proliferation through a multitarget effect.

Overall, the physical-chemical features and antimicrobial activity of IPFs make them good formulations to develop green, bio-compatible, and efficient coatings that can prevent pathogen growth and limit AMR phenomena.

## Author contributions

**Elena Piacenza:** Conceptualization, Methodology, Validation, Formal analysis, Investigation, Data curation, Writing - Original draft, Visualization. **Alessandro Presentato:** Conceptualization, Methodology, Validation, Formal analysis, Data curation, Writing - Original Draft. **Rosa Alduina:** Resources, Writing - Review & Editing, Supervision. **Antonino Scurria:** Methodology, Writing - Review & Editing. **Mario Pagliaro:** Resources, Writing - Review & Editing, Supervision, Project administration. **Lorenzo Albanese:** Resources. **Francesco Meneguzzo:** Resources, Writing - Review & Editing. **Rosaria Ciriminna:** Resources, Writing - Review & Editing, Supervision. **Delia F. Chillura Martino:** Resources, Writing - Review & Editing, Supervision, Project administration, Funding acquisition.

## Acknowledgments

We acknowledge the Italian Ministry of University and Research (MUR) for the PON project on Research and Innovation 2014-2020 (Azione IV.4 - Contratti di ricerca su tematiche dell’Innovazione - B75F21002190001) and the PON Project on Research and Innovation 2012–2020 (Attraction and International Mobility – AIM1808223). Antonio Scurria gratefully acknowledges the MUR and the Università degli Studi di Messina for funding his PhD program (D. M. 1061/2021). We also thank Campisi Citrus (Siracusa, Italy) for the gift of waste lemon and grapefruit peel from which IntegroPectins were obtained, and ATeN Center (Palermo, Italy) for the access to NMR and XRD instrumentation.

---

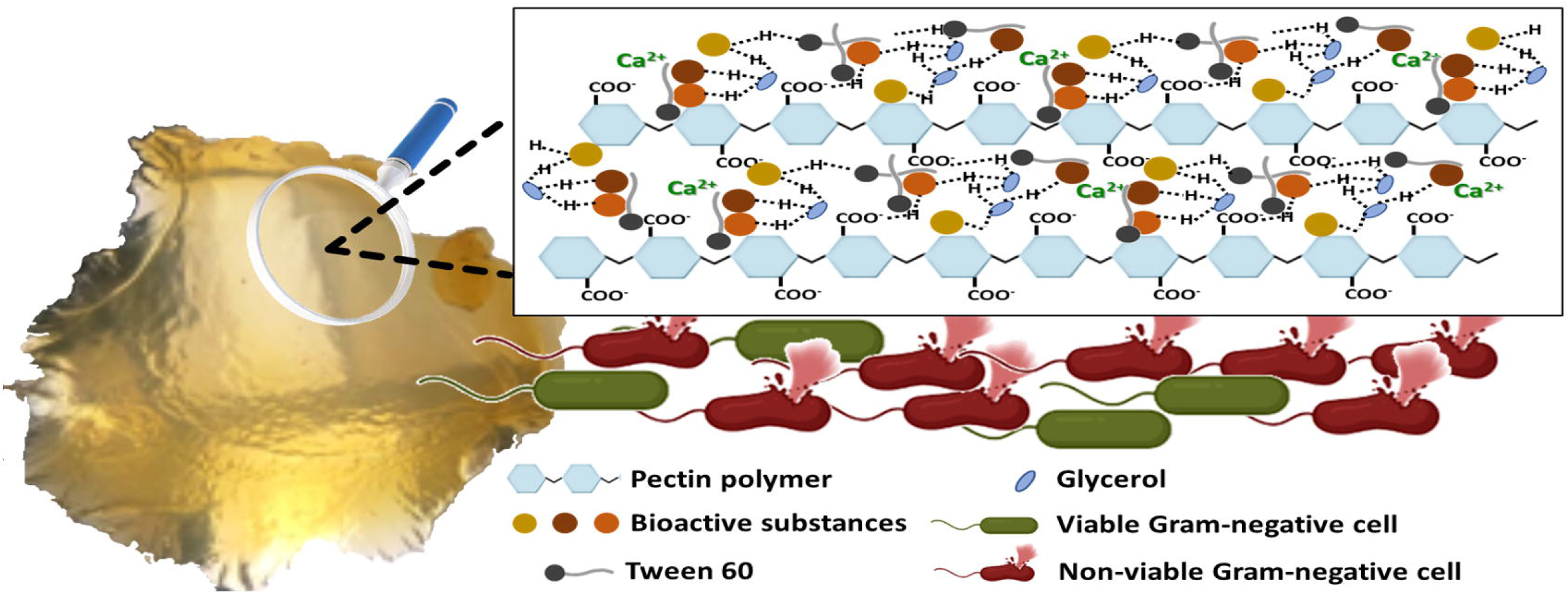

