## Supplementary Information for "Cross-linked natural IntegroPectin films from Citrus biowaste with intrinsic antimicrobial activity"

[a]*Department of Biological, Chemical, and Pharmaceutical Sciences and Technologies (STEBICEF), University of Palermo, viale delle Scienze, 90128 Palermo (PA), Italy;* [b]*Istituto per lo Studio dei Materiali Nanostrutturati, CNR, via U. La Malfa 153, 90146 Palermo (PA), Italy;* [c]*Istituto per la Bioeconomia, CNR, via Madonna del Piano 10, 50019 Sesto Fiorentino (FI), Italy;* [d]*National Interuniversity Consortium of Materials Science and Technology (INSTM), via G. Giusti 9, 50121 Firenze (FI), Italy*

### *Supplementary Information*

**Corresponding authors*:
Dr. E. Piacenza
Department of Biological, Chemical, and Pharmaceutical Sciences and Technologies (STEBICEF)
University of Palermo
viale delle Scienze, 90128 Palermo (Italy)


Dr. R. Ciriminna
Istituto per lo Studio dei Materiali Nanostrutturati, CNR
via U. La Malfa 153, 90146 Palermo (Italy)


**Table S1.** ATR-FTIR absorption bands of films obtained from commercial citrus pectin (CPF), lemon IntegroPectin (LIPF), and grapefruit IntegroPectin (GIPF).

| $\tilde{\nu}$ [cm$^{-1}$][a] | | | Vibrational mode | Identification |
|---|---|---|---|---|
| CPF | LIPF | GIPF | | |
| 3316 | | | $\nu^{b}$ (OH) | Polysaccharides (Synytsya et al., 2003a; Fidalgo et al., 2016); water (Synytsya et al., 2003a; Fidalgo et al., 2016) |
| | 3292 | 3293 | $\nu$ (OH) | Polysaccharides (Synytsya et al., 2003a; Fidalgo et al., 2016); water (Synytsya et al., 2003a; Fidalgo et al., 2016); polyphenols (Heredia-Guerrero et al., 2014); glycerol (Danish et al., 2017) |
| 2926 | 2930 | 2931 | $\nu_{as}^{c}$ ($CH_2$); $\nu_{as}$ ($CH_3$); $\nu$ (CH) | Pectin backbone (Fidalgo et al., 2016); arabinose and galactose (Fidalgo et al., 2016); glycerol (Danish et al., 2017); Tween 60 (Branzoi & Branzoi, 2017) |
| | 2880 | 2882 | $\nu_{s}^{d}$ ($CH_3$); $\nu$ (CH) | Pectin backbone (Fidalgo et al., 2016); pyranose rings (Fidalgo et al.,2016); glycerol (Danish et al., 2017) |
| | 2854 | 2856 | $\nu$ (CH) | Pectin backbone (Bichara et al., 2016); glycerol (Danish et al., 2017); Tween 60 (Branzoi & Branzoi, 2017) |
| | 2657 | 2658(s) | $\nu$ (OH) | Free carboxylic acids (Synytsya et al., 2003a; Fidalgo et al., 2016) |
| 1743 | | | $\nu$ $(C{=}O)_{ester}$ | Methyl esterified carboxylic groups of galacturonic acid (Synytsya et al., 2003a; Fidalgo et al., 2016; Manrique & Lajolo, 2002; Kyomugasho et al., 2015; Wang et al., 2016; La Cava et al., 2018) |
| | 1717 | 1718 | $\nu_{s}$ (C=O) | Carboxylate and nonconjugated keto groups of carotenoids (Synytsya et al., 2003a; Fidalgo et al., 2016; Aburto et al., 2015; Bichara et al., 2016), phenols and flavonoids (Heredia-Guerrero et al., 2014; Lóránd et al., 2002; Ricci et al., 2015); terpenes (Schulz & Baranska 2007) |
| | | 1640(s) | $\delta^{e}$ ($H_2O$); $\nu$ (C=C); $\nu$ (C-C); $\nu$ (C=C) | Phenyl and uracyl groups (La Cava et al., 2018); phenolic acids (Heredia-Guerrero et al., 2014; Ricci et al., 2015); alkenes of essential oils (Cebi et al., 2021); terpenes (Schulz & Baranska 2007) |
| 1611 | 1604 | 1604 | $\nu_{as}$ ($COO^-$); $\nu$ (C-C); benzene ring | Carboxylate groups of polygalacturonic acid (Fidalgo et al., 2016; La Cava et al., 2018; Aburto et al., 2015; Bichara et al., 2016); polyphenols (Schulz & Baranska 2007) |
| | 1514 | 1516 | $\delta_{ip}^{f}$ (CH); $\nu$ (C=C) | Phenyl rings (Heredia-Guerrero et al., 2014; Ricci et al., 2015; Zeier & Schreiber, 1999); carotenoid compounds (Lóránd et al., 2002) |
| | 1460(s) | 1459(s) | $\delta_{as}$ ($CH_3$); $\delta$ ($CH_2$); | Pectin backbone (Bichara et al., 2016; Zeier & Schreiber, 1999; Rashid et al., 2016; Sene et al., |

| $\tilde{\nu}$ [cm$^{-1}$][a] | | | Vibrational mode | Identification |
|---|---|---|---|---|
| | | | ν (C=C);<br>$\nu_{as}$ ($COO^-$);<br>$\beta^g$ (furanic ring) | 1994); aromatic compounds (Ricci et al., 2015; Suresh et al., 2018; Tracanna et al., 2019; Schulz & Baranska, 2009); terpenes (Schulz & Baranska 2007); Tween 60 (Branzoi & Branzoi, 2017); glycerol (Kataoka et al., 2011) |
| 1439(s) | 1440(s) | 1438(s) | $\delta_{as}$ ($CH_3$);<br>$\rho^h$ ($CH_2$) 1st overtone;<br>ν (C-C);<br>δ (CH); ρ (CH) | Aromatic compounds (Heredia-Guerrero et al., 2014; Ricci et al., 2015; Sene et al., 1994; Suresh et al., 2018; Tracanna et al., 2019); ester methyl groups in galacturonic and rhamnose rings of pectin (Synytsya et al., 2003a; Fidalgo et al., 2016; Aburto et al., 2015; Bichara et al., 2016; Schulz & Baranska, 2009); alkanes of terpenes (Cebi et al., 2021); polygalacturonic acid (Bichara et al., 2016) |
| 1413 | 1410 | 1412 | δ $(COH)_{COOH}$;<br>$\nu_s$ ($COO^-$);<br>ρ (CH);<br>$\delta_{ip}$ (CH);<br>$\delta_s$ ($CH_3$);<br>δ (COH) | Methyl groups (Heredia-Guerrero et al., 2014; Manrique & Lajolo, 2002); ring vibration (Heredia-Guerrero et al., 2014; Bichara et al., 2016; Ricci et al., 2015); carboxylate pectin ester groups (Synytsya et al., 2003a; Heredia-Guerrero et al., 2014; Manrique & Lajolo, 2002); polygalacturonic acid (Bichara et al., 2016); glycerol (Danish et al., 2017; Kataoka et al., 2011) |
| 1368 | 1367 | 1369 | $\delta_s$ ($CH_2$);<br>$\delta_s$ ($CH_3$);<br>$\beta_s$ ($CH_3$);<br>δ (OH);<br>$\delta_{ip}$ (COH);<br>$\delta_{op}^i$ ($CH_3$) | Ester methyl groups in galacturonic and rhamnose rings of pectin (Synytsya et al., 2003a; Fidalgo et al., 2016; Bichara et al., 2016; Sene et al., 1994); flavonoids (Ricci et al., 2015; Suresh et al., 2018; Tracanna et al., 2019; Schulz & Baranska, 2009) |
| | 1352 | 1354 | ρ (CH) | Polygalacturonic acid (Bichara et al., 2016); Tween 60 (Branzoi & Branzoi, 2017) |
| 1330 | 1329 | 1331 | δ (CH);<br>$\nu_s$ ($COO^-$);<br>$\omega^j$ ($CH_2$);<br>$\delta_{ip}$ (C-O-H) | Pyranose in pectic ring (Synytsya et al., 2003a; Ricci et al., 2015; Copikova et al., 2001); metoxyphenolic substitutions (Bichara et al., 2016); alcohol hydroxyl groups in pyranose ring (Fidalgo et al., 2016) |
| 1264 | | | β (OH);<br>ν (C-O-C) | Hydroxyl groups of polysaccharides (Fidalgo et al., 2016; Heredia-Guerrero et al., 2014; Bichara et al., 2016; Sene et al., 1994; Schulz & Baranska, 2009; Copikova et al., 2001) |
| | | 1250(s) | ν (C-O);<br>δ (CH); δ (OH) | Polyols (hydroxyflavonoids) (Sene et al., 1994; Suresh et al., 2018; Tracanna et al., 2019; Schulz & Baranska, 2009); polygalacturonic acid (Bichara et al., 2016) |
| 1231 | 1226 | 1225 | ν ($C(CH_3)_2$);<br>ν (CO-O);<br>δ (OH);<br>$\nu_{ip}$ (OH); | Aromatic ethers (Heredia-Guerrero et al., 2014; Lóránd et al., 2002; Ricci et al., 2015; Suresh et al., 2018; Tracanna et al., 2019; Schulz & Baranska, 2009; Copikova et al., 2001); metoxyphenolic substitutions (Lóránd et al., |

| $\tilde{\nu}$ [cm$^{-1}$][a] | | | Vibrational mode | Identification |
|---|---|---|---|---|
| | | | $\delta_{ip}$ (C-O-H); | 2002); alcohol hydroxyl groups in pyranose ring (Synytsya et al., 2003a; Fidalgo et al., 2016); carboxylic groups of polygalacturonic acid (Bichara et al., 2016) |
| | 1199(s) | 1198(s) | ν (C(CH$_3$)$_2$); ν (C-C) | cyclic C-C bonds in the pectin ring (Synytsya et al., 2003a; Fidalgo et al., 2016; Manrique & Lajolo, 2002; Kyomugasho et al., 2015; Aburto et al., 2015; Bichara et al., 2016; Sene et al., 1994); flavonoids (Heredia-Guerrero et al., 2014; Sene et al., 1994; Suresh et al., 2018; Tracanna et al., 2019; Schulz & Baranska, 2009) |
| 1142 | 1141 | 1140 | ν (C-O-C); ν (C-C); $\nu_{as}$ (O-C-O) | Glycosidic bond in polysaccharide ring (Synytsya et al., 2003; Fidalgo et al., 2016; Aburto et al., 2015; Bichara et al., 2016; Copikova et al., 2001); cyclic C-C bonds in the pectin ring (La Cava et al., 2018; Bichara et al., 2016) |
| 1102 | 1100 | 1100 | ν (C-O); ν (C-OH); ν (C-O-C); ν (C-C) | Pyranose and glycoside (Synytsya et al., 2003a; Fidalgo et al., 2016; Kyomugasho et al., 2015; Copikova et al., 2001); pectin ring (Synytsya et al., 2003a; Aburto et al., 2015; Bichara et al., 2016; Schulz & Baranska, 2009); uronic acid (Heredia-Guerrero et al., 2014; La Cava et al., 2018); glycerol (Kataoka et al., 2011) |
| 1096 | | | ν (C-O); ν (C-OH); ν (C-O-C); ν (C-C) | Pyranose and glycoside (Synytsya et al., 2003a; Fidalgo et al., 2016; Zeier & Schreiber, 1999; Copikova et al., 2001); pectin ring (Synytsya et al., 2003a; Aburto et al., 2015; Bichara et al., 2016; Schulz & Baranska, 2009); uronic acid (Heredia-Guerrero et al., 2014; La Cava et al., 2018) |
| 1081 | | | ν (C-O); δ (OH) | Polygalacturonic acid (Bichara et al., 2016) |
| 1073 | | | ν (C-O); ρ (CO); ν (C-O-C); ν (C-C) ν (C-OH) | Pyranose and glycoside (Synytsya et al., 2003a; Fidalgo et al., 2016; Bichara et al., 2016; Schulz & Baranska, 2009); arabinose and galactose (La Cava et al., 2018) |
| 1046 | | | ν (C-O); ρ (CO); ν (C-O-C); ρ (CH$_3$) ν (C-OH); ν (C-C) | Pyranose and glycoside (Synytsya et al., 2003a; Fidalgo et al., 2016; Aburto et al., 2015; Bichara et al., 2016; Schulz & Baranska, 2009); arabinose and galactose (La Cava et al., 2018) |
| | 1034 | 1032 | ν (C-O); ν (C-O-C); ν (COH) | Esterified polygalacturonic acid (Bichara et al., 2016); terpenes (Schulz & Baranska 2007); glycerol (Adamu et al., 2017); Tween 60 (Branzoi & Branzoi, 2017) |
| 1018 | | | ν (C-O) | Polygalacturonic acid (Bichara et al., 2016) |

| $\tilde{\nu}$ [cm$^{-1}$][a] | | | Vibrational mode | Identification |
|---|---|---|---|---|
| 1011 | | | ν (C-C); ν (C-O) | Polysaccharides (Synytsya et al., 2003a; Heredia-Guerrero et al., 2014; Copikova et al., 2001); pectin (C2-C3, C2-O2, C1-O1) (Bichara et al., 2016; Schulz & Baranska, 2009); uronic acid (La Cava et al., 2018) |
| | 991 | 992 | ν (C-C); ν (C-O); δ ($COO^-$); β (CC) | Polygalacturonic acid (Bichara et al., 2016); terpenes (Schulz & Baranska 2007); glycerol (Kataoka et al., 2011; Danish et al., 2017) |
| | 970(s) | 967(s) | $\gamma^k$ (=CH); ρ ($CH_3$); ν$(C=C)_{trans}$; ν (C-O) | Polysaccharides (Synytsya et al., 2003a; Fidalgo et al., 2016; Heredia-Guerrero et al., 2014; La Cava et al., 2018; Copikova et al., 2001); arabinose and galactose (La Cava et al., 2018); flavonoids (Heredia-Guerrero et al., 2014; Ricci et al., 2015; Zeier & Schreiber, 1999; Sene et al., 1994; Suresh et al., 2018; Tracanna et al., 2019; Schulz & Baranska, 2009) |
| 954 | | | δ (C=O); δ (CCH); δ (COH) | Polysaccharides in pectin (Synytsya et al., 2003a; Fidalgo et al., 2016; Heredia-Guerrero et al., 2014; La Cava et al., 2018; Copikova et al., 2001) |
| 915 | 922 | 921 | ρ ($CH_3$); α-anomeric linkage; $\delta_{op}$ $(=CH)_{trans}$; β (Ph); $\tau^l$ (HCC) | Ester methyl groups (Fidalgo et al., 2016; Bichara et al., 2016; Copikova et al., 2001); glucose and fructose (Synytsya et al., 2003a; Fidalgo et al., 2016; Bichara et al., 2016); phenyl moieties (Heredia-Guerrero et al., 2014; Ricci et al., 2015; Zeier & Schreiber, 1999; Sene et al., 1994; Suresh et al., 2018; Tracanna et al., 2019); pectin (Synytsya et al., 2003a; Wang et al., 2016; Schulz & Baranska, 2009); flavonoids (Heredia-Guerrero et al., 2014; Ricci et al., 2015; Zeier & Schreiber, 1999; Sene et al., 1994; Suresh et al., 2018; Tracanna et al., 2019; Schulz & Baranska, 2009) |
| 890 | | | ν (C-C); $\delta_{op}$ (CH); δ (CCH); δ (COH) | Pectin (Synytsya et al., 2003a; Fidalgo et al., 2016; Heredia-Guerrero et al., 2014; Wang et al., 2016; Schulz & Baranska, 2009) |
| 880 | | 884(s) | β (CH); δ (CCH); δ (COH); γ (=CH); $\delta_{op}$ ($C=CH_2$); ν (C-O) | Methylene groups (Fidalgo et al., 2016; Bichara et al., 2016; Copikova et al., 2001); vinilydiene groups of terpenoids (Mitzner & Theimer, 1962); polygalacturonic acid (Bichara et al., 2016) |
| | 865(s) | 866(s) | ρ ($CH_2$); $\delta_{ip}$ (CH); ρ ($CH_2$); β ($C-C_{ring}$) | Pyranose (Synytsya et al., 2003a; Fidalgo et al., 2016; Aburto et al., 2015; Bichara et al., 2016; Schulz & Baranska, 2009); phenols (Heredia-Guerrero et al., 2014; Ricci et al., 2015; Zeier & Schreiber, 1999; Sene et al., 1994; Suresh et al., 2018; Tracanna et al., 2019; Schulz & Baranska, |

| $\tilde{\nu}$ [cm$^{-1}$][a] | | | Vibrational mode | Identification |
|---|---|---|---|---|
| | | | | 2009) |
| 850 | 848 | 849 | ρ ($CH_2$); β (CC) | Glycerol (Danish et al., 2017) |
| 833 | 832 | 833 | γ (OH); $\delta_{op}$ (CH) | Six-membered ring of polyphenols (Heredia-Guerrero et al., 2014; Ricci et al., 2015; Zeier & Schreiber, 1999; Sene et al., 1994; Suresh et al., 2018; Tracanna et al., 2019; Schulz & Baranska, 2009); α-glycosidic linkages (La Cava et al., 2018); phenolic compounds (Heredia-Guerrero et al., 2014; Ricci et al., 2015; Zeier & Schreiber, 1999; Sene et al., 1994; Suresh et al., 2018; Tracanna et al., 2019; Schulz & Baranska, 2009) |
| 809(s) | 815(s) | 812 | ρ ($CH_2$); $\delta_{ip}$ (C-H); $\beta_{op}$ (CH); $\omega_{op}$ (CH) | Pyranose (Synytsya et al., 2003a; Fidalgo et al., 2016; Aburto et al., 2015; Bichara et al., 2016; Schulz & Baranska, 2009); polyphenols (Schulz & Baranska 2007) |
| 783 | | 779(s) | ω (CH); ρ ($CH_2$); $\delta_{ip}$ (C-H); γ (COH); $\delta_{op}$ (=CH) *cis* | Pyranose (Synytsya et al., 2003a; Fidalgo et al., 2016; Aburto et al., 2015; Bichara et al., 2016; Schulz & Baranska, 2009); six-membered ring of polyphenols (Heredia-Guerrero et al., 2014; Ricci et al., 2015; Zeier & Schreiber, 1999; Sene et al., 1994; Suresh et al., 2018; Tracanna et al., 2019; Schulz & Baranska, 2009); polygalacturonic acid (Bichara et al., 2016) |
| 771 | | | δ ($COO^-$); ring breezing | Esterified polygalacturonic acid (Bichara et al., 2016) |
| 758 | 759 | 759(s) | $\delta_{op}$ $(=CH)_{cis}$; breath[m] | Breathing ring (Fidalgo et al., 2016; Bichara et al., 2016) |
| 746 | | 744(s) | γ $(COH)_{COOH}$ | Pectin (Bichara et al., 2016) |
| 729 | 730(s) | 734(s) | δ (OCC) | Polygalacturonic acid (Bichara et al., 2016) |
| 715 | 714 | 716 | ρ ($CH_2$); γ (COH); $\delta_{op}$ (=CH); $\delta_{op}$ (=CH)cis | six-membered ring of polyphenols (Heredia-Guerrero et al., 2014; Ricci et al., 2015; Zeier & Schreiber, 1999; Sene et al., 1994; Suresh et al., 2018; Tracanna et al., 2019; Schulz & Baranska, 2009); vibrations of pyranoid ring (Synytsya et al., 2003a; Fidalgo et al., 2016; Aburto et al., 2015; Bichara et al., 2016; Schulz & Baranska, 2009) |
| | | 703 | $\delta_{op}$ (=CH); γ $(COH)_{ring}$ | Pectin (Synytsya et al., 2003a; Fidalgo et al., 2016; Heredia-Guerrero et al., 2014; Wang et al., 2016; Schulz & Baranska, 2009; Bichara et al., 2016) |
| 683 | | | ω (C=O); $\delta_{op}$ (=CH); $\nu_s$ (C-O-C) | Glycoside linkage (Synytsya et al., 2003a; Fidalgo et al., 2016); acidic pectins (Synytsya et al., 2003a; Fidalgo et al., 2016; Heredia-Guerrero et al., 2014; Wang et al., 2016; Schulz & Baranska, 2009) |
| | 666 | 665 | β (C-C-O); | Phenols (Heredia-Guerrero et al., 2014; Ricci et |

| $\tilde{\nu}$ [cm$^{-1}$][a] | Vibrational mode | Identification |
|---|---|---|
| | γ (C-O) | al., 2015; Zeier & Schreiber, 1999; Sene et al., 1994; Suresh et al., 2018; Tracanna et al., 2019; Schulz & Baranska, 2009) |

[a] $\tilde{\nu}$ [cm$^{-1}$] = wavenumber; [b] ν = stretching vibration; [c] as = asymmetric vibration; [d] s = symmetric vibration; [e] δ = bending/scissoring vibration; [f] ip = in plane vibration; [g] β = deformation modes; [h] ρ = rocking vibration; [i] op = out of plane vibration; [j] ω = wagging vibration; [k] γ = out of plane ring vibrations; [l] τ = twisting vibration; [m] breath = breathing mode.

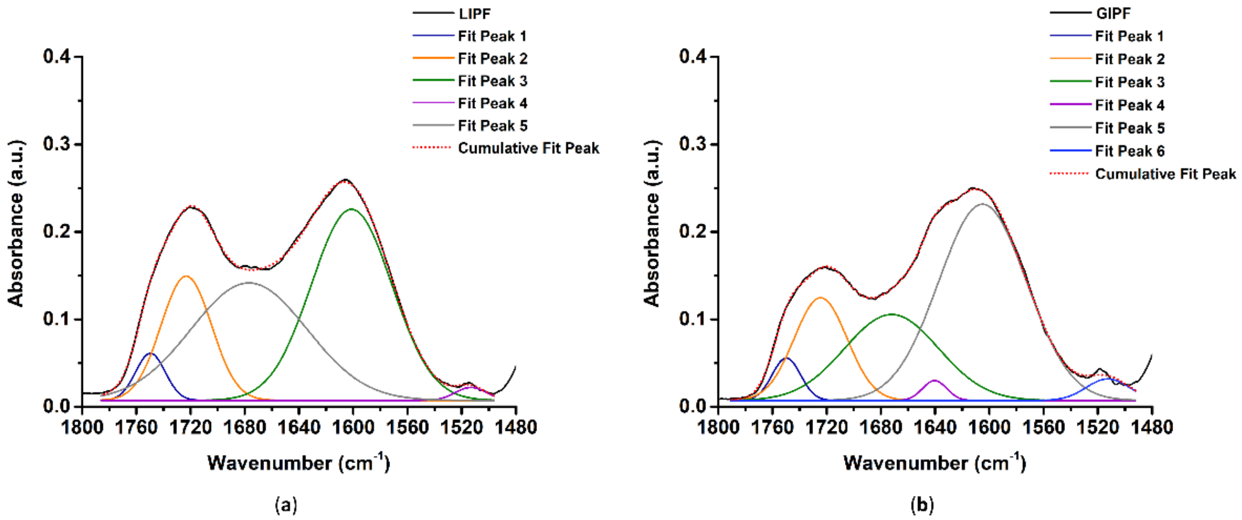


**Figure S1.** ATR-FTIR spectral deconvolution by non-linear least-squares fitting of LIPF (**a**) and GIPF (**b**) in the 1800-1480 cm$^{-1}$ region.

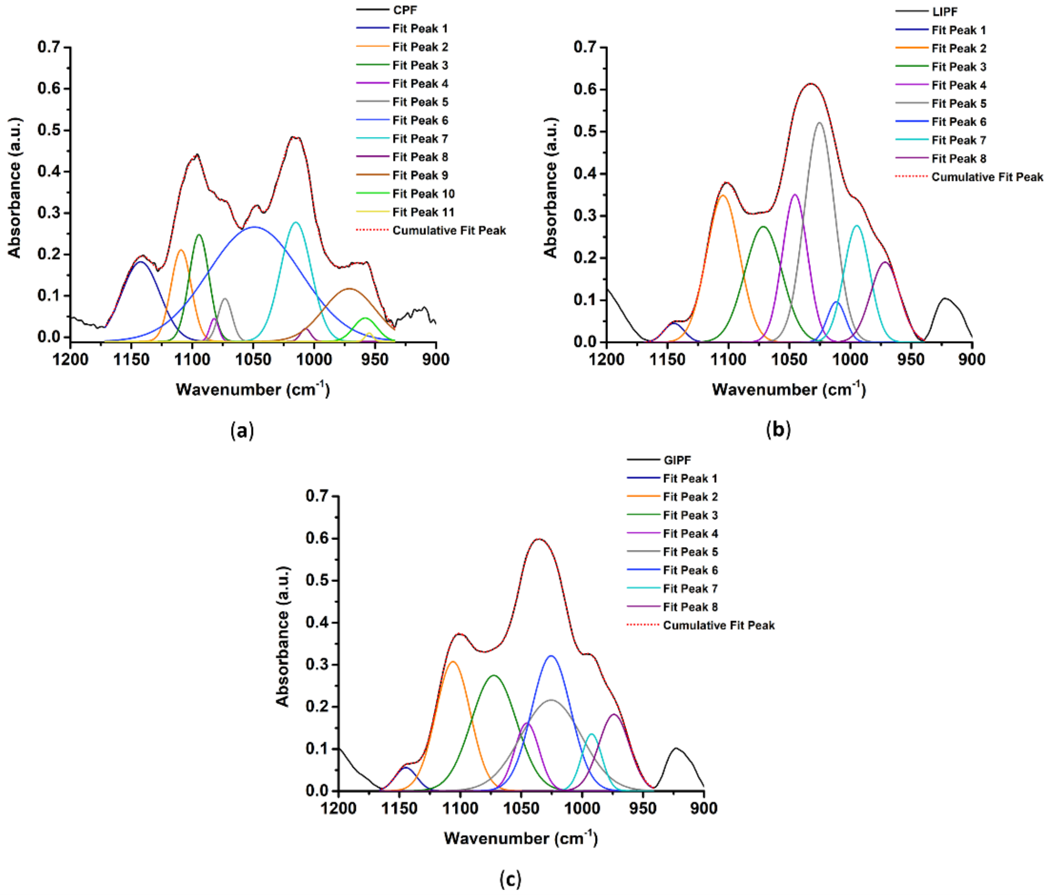


**Figure S2.** ATR-FTIR spectral deconvolution by non-linear least-squares fitting of CPF (a), LIPF (b) and GIPF (c) in the 1200-900 $cm^{-1}$ region.

**Table S2.** $^{13}C$ CPMAS NMR chemical shifts and their identification for CPF, LIPF, and GIPF.

| $\sigma$ [ppm][a] | | | Resonance mode | Identification |
|---|---|---|---|---|
| CPF | LIPF | GIPF | | |
| 177.11 | | | $COO^-$ (C6) | Galacturonic units (Gal) (Synytsya et al., 2003b) |
| 171.50 | 171.07 | 171.16 | O<u>C</u>OCH$_3$ (C6) | Acetyl groups in Gal (Synytsya et al., 2003b) |
| 101.17 | 101.21 | 100.42 | C1 | Glycosidic bond of pectin (Synytsya et al., 2003b; Soltys et al., 2019) |
| | 100.3 | 99.00 | C1 | Glycosidic bond of pectin (Synytsya et al., 2003b; Soltys et al., 2019) |
| 80.11 | 81.04 | 80.65 | C4 | Glycosidic bond of pectin (Synytsya et al., 2003b) |
| 77.42 | | 77.49 | | |
| 73.02 | 73.05 | | C5, C3, C2 | Pyranoid ring (Synytsya et al., 2003b) |
| 70.91 | 70.80 | | | |
| 69.00 | 69.61 | 70.00 | | |
| 63.60 | 63.63 | 63.68 | C2 | Glycerol (Soltys et al., 2019) |
| 53.61 | 53.66 | 53.54 | –COO<u>C</u>H$_3$ | Methyl esters in pectin (Synytsya et al., 2003b) |
| | 45.06 | 43.36 | $CR_4$ | Mixture of terpenoids (Conte et al., 2010) |
| 33.34 | | 33.11 | $(–(CH_2)_n–)$ | Aliphatic chains of Tween 60 (Zhang et al., 2015) |
| 30.90 | 30.46 | 30.47 | $(–(CH_2)_n–)$ | Aliphatic chains of Tween 60 (Zhang et al., 2015) |

[a)] $\sigma$ [ppm] = chemical shift.

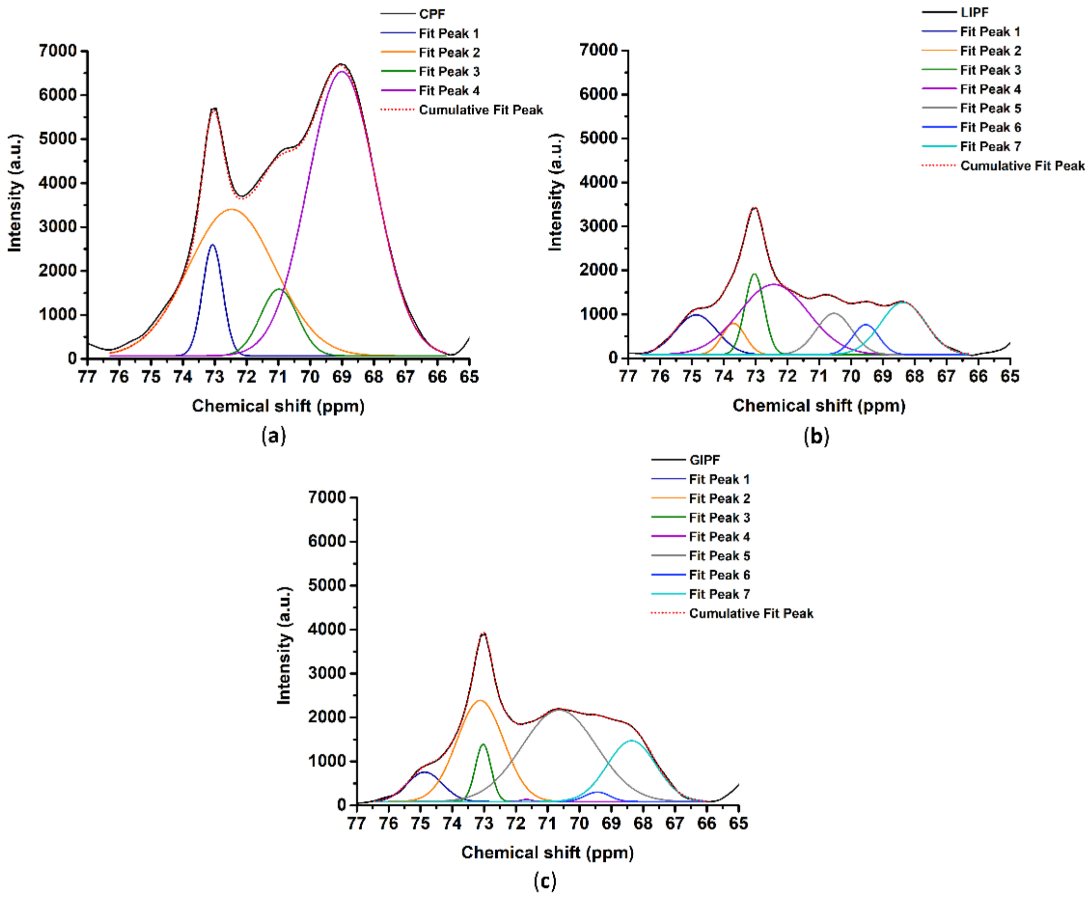


**Figure S3.** $^{13}C$ CPMAS NMR spectral deconvolution by non-linear least-squares fitting of CPF (**a**), LIPF (**b**) and GIPF (**c**) in the 76-66 ppm region.

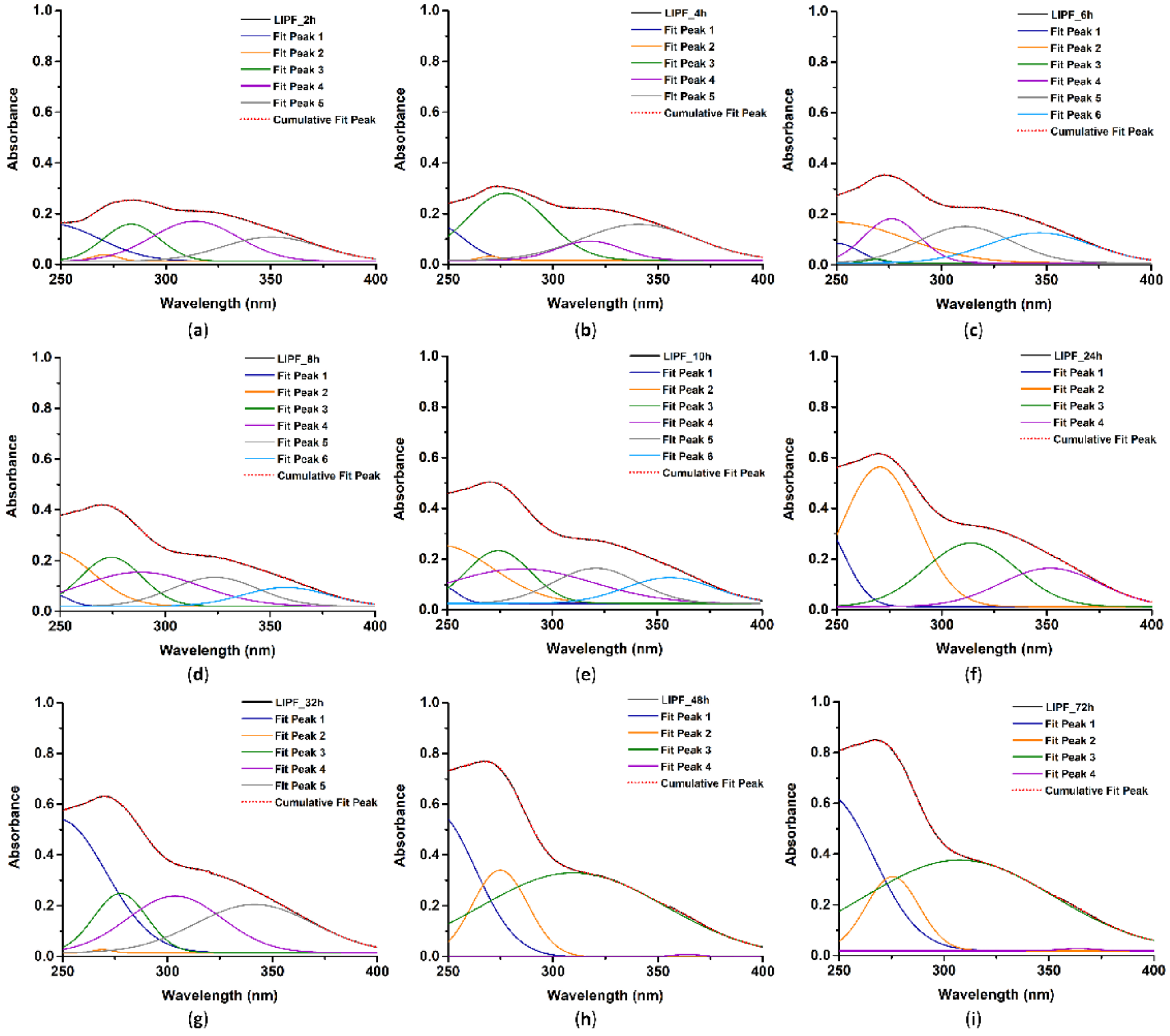


**Figure S4.** UV-Visible spectral deconvolution in the 250-400 nm region by non-linear least-squares fitting of LIPF incubated in LB medium for 2 (a), 4 (b), 6 (c), 8 (d), 10 (e), 24 (f), 32 (g), 48 (h), and 72h (i).

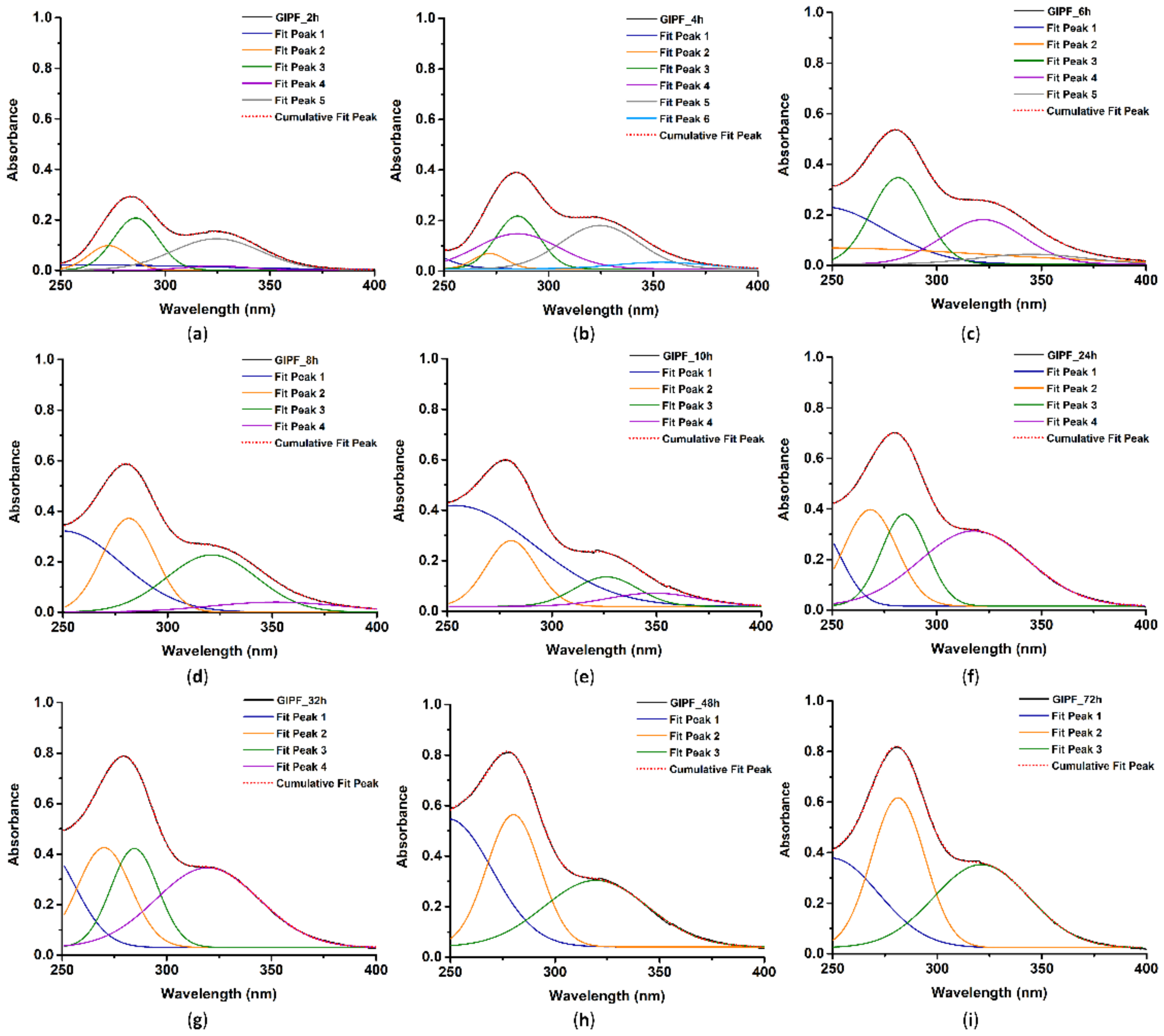


**Figure S5.** UV-Visible spectral deconvolution in the 250-400 nm region by non-linear least-squares fitting of GIPF incubated in LB medium for 2 (**a**), 4 (**b**), 6 (**c**), 8 (**d**), 10 (**e**), 24 (**f**), 32 (**g**), 48 (**h**), and 72h (**i**).

**Table S3.** Deconvolution of UV-Visible spectra of IPFs in the 250-400 nm region recorded in the 2-72h timeframe.

| Samples | $\lambda^a$ [nm] | $w^b$ | $A^c$ | Samples | λ [nm] | w | A |
|---|---|---|---|---|---|---|---|
| LIPF_2h | 271 | 12.81 | 0.420 | GIPF_2h | 273 | 18.98 | 2.308 |
| | 283 | 26.72 | 4.911 | | 286 | 21.22 | 5.4870 |
| | 314 | 41.14 | 8.143 | | 324 | 42.11 | 7.554 |
| | 350 | 46.30 | 5.583 | GIPF_4h | 272 | 59.71 | 1.197 |
| LIPF_4h | 270 | 9.703 | 0.200 | | 285 | 20.60 | 12.32 |
| | 278 | 38.46 | 12.77 | | 324 | 37.05 | 7.977 |
| | 318 | 31.09 | 2.978 | | 354 | 43.59 | 1.430 |
| | 341 | 53.99 | 9.630 | GIPF_6h | 281 | 25.93 | 11.13 |
| LIPF_6h | 269 | 9.895 | 0.185 | | 322 | 39.19 | 8.662 |
| | 276 | 26.12 | 5.712 | | 345 | 51.38 | 2.550 |
| | 311 | 43.96 | 7.933 | GIPF_8h | 282 | 25.36 | 11.80 |
| | 347 | 49.68 | 7.418 | | 321 | 43.02 | 63.21 |
| LIPF_8h | 274 | 29.07 | 7.039 | | 352 | 63.21 | 3.161 |
| | 288 | 56.66 | 9.570 | GIPF_10h | 281 | 23.91 | 7.800 |
| | 324 | 41.08 | 5.902 | | 326 | 30.17 | 4.417 |
| | 358 | 40.76 | 3.804 | | 349 | 43.28 | 2.854 |
| LIPF_10h | 274 | 30.56 | 7.985 | GIPF_24h | 270 | 25.28 | 12.11 |
| | 285 | 68.41 | 11.72 | | 285 | 21.94 | 10.00 |
| | 321 | 40.06 | 6.989 | | 318 | 51.73 | 19.33 |
| | 356 | 42.38 | 5.431 | GIPF_32h | 270 | 25.58 | 12.67 |
| LIPF_24h | 269 | 9.198 | 0.143 | | 285 | 22.48 | 11.04 |
| | 278 | 25.21 | 7.374 | | 320 | 49.30 | 19.50 |
| | 304 | 45.12 | 12.64 | GIPF_48h | 280 | 25.51 | 14.10 |
| | 342 | 56.15 | 13.40 | | 325 | 35.26 | 6.104 |
| LIPF_32h | 270 | 35.55 | 24.55 | GIPF_72h | 281 | 25.52 | 18.98 |
| | 314 | 43.28 | 13.60 | | 321 | 45.52 | 18.77 |
| | 352 | 47.05 | 8.958 | | | | |
| LIPF_48h | 275 | 26.47 | 11.35 | | | | |
| | 310 | 87.43 | 36.34 | | | | |
| | 364 | 15.09 | 0.190 | | | | |
| LIPF_72h | 275 | 25.11 | 9.193 | | | | |
| | 308 | 88.92 | 39.82 | | | | |
| | 367 | 16.49 | 0.189 | | | | |

[a] λ [nm] = wavelength; [b] w = width; [c] A = integrated area.

**Table S4.** Release parameters obtained by using the first-order kinetic, second-order kinetic, and Ritger-Peppas models on UV-visible cumulative data.

| Absorbance contributions | First-order kinetic | | Second-order kinetic | | Ritger-Peppas model | | |
|---|---|---|---|---|---|---|---|
| | $R^{2\,a}$ | $k^{b}$ | $R^2$ | k | $R^2$ | $n^{c}$ | k |
| LIPF_$A_{270-275}$ | 0.9140 | 0.083 | 0.9654 | 0.1406 | 0.9789 | 0.3297 | 0.2496 |
| LIPF_$A_{306-315}$ | 0.8101 | 0.1511 | 0.8071 | 0.2781 | 0.9468 | 0.2001 | 0.4248 |
| LIPF_$A_{340-350}$ | 0.8427 | 0.1979 | 0.8253 | 0.3913 | 0.9343 | 0.1916 | 0.4757 |
| GIPF_$A_{280-285}$ | 0.9493 | 0.1553 | 0.9680 | 0.2727 | 0.9145 | 0.2361 | 0.3891 |

[a)] $R^2$ = R-squared value; [b)] k = rate constant; [c)] n = release exponent.

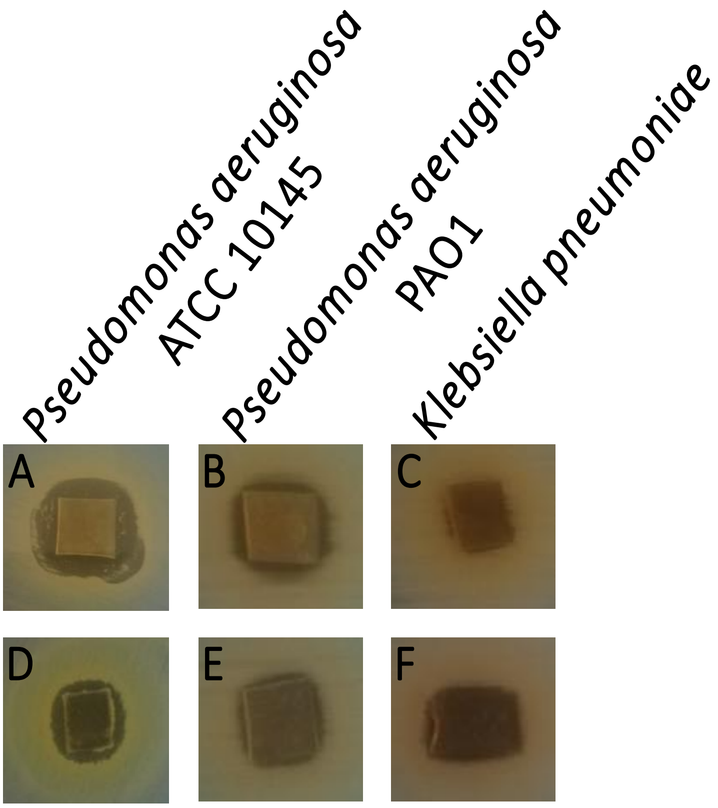


**Figure S6.** Antimicrobial activity of LIPF (**a, b,** and **c**) and GIPF **(d, e,** and **f**) against adherent growing *Pseudomonas aeruginosa* ATCC 10145, *Pseudomonas aeruginosa* PAO1, and *Klebsiella pneumoniae* cells.
